# Social Decision Preferences for Close Others are Embedded in Neural and Linguistic Representations

**DOI:** 10.1101/2024.07.16.603808

**Authors:** João F. Guassi Moreira, L. Concepción Esparza, Jennifer A. Silvers, Carolyn Parkinson

## Abstract

Humans frequently make decisions that impact close others. Prior research has shown that people have stable preferences regarding such decisions and maintain rich, nuanced mental representations of their close social partners. Yet, if and how such mental representations shape social decisions preferences remains to be seen. Using a combination of functional magnetic resonance imaging (fMRI) and natural language processing (NLP), this study investigated how neural and linguistic representations of close others relate to social decision-making. After nominating a parent and friend, male and female participants (*N* = 63) rated their characteristics and made hypothetical social decisions while undergoing fMRI. Neural representations of parents and friends, relative to the self, were linked to social decision preferences. Specifically, greater neural similarity between self and parent in the temporoparietal junction (TPJ) and nucleus accumbens (NAcc) was associated with a preference for parents, while greater self-friend similarity in the medial prefrontal cortex (mPFC) was associated with friend-preference. Additionally, linguistic analysis of written descriptions of close others from a separate sample of males and females (*N* = 1,641) revealed that social decision preferences could be reliably predicted from semantic features of the text. High correspondence between neural and linguistic data in the imaging sample further strengthened the association with social decision preferences. These findings help elucidate the neural and linguistic underpinnings of social decision-making, emphasizing the critical role of mental representations in guiding choices involving familiar others.

**Significance Statement:** This study provides novel insights into how mental representations of close others relate to social decision-making. By combining brain imaging and natural language processing, we show that both neural and linguistic representations of familiar individuals (parents and friends), can predict social preferences. We found that neural representations of these close others are linked to the choices people make about these individuals. Additionally, the way people describe their close others in writing are reliably associated with their decision preferences. Our approach, integrating neuroscience and language analysis, significantly advances our understanding of the cognitive mechanisms behind social decision-making, posing implications for fields ranging from psychology to artificial intelligence. These findings highlight the complexity of human relationships and their impact on everyday decisions.

## Introduction

Humans are frequently faced with decisions that could impact multiple other people. For instance, one may need to choose between having dinner with a close friend or attending happy hour with a new co-worker, or whether to buy a plane ticket home for the holidays to see family or use that money to take a trip abroad with a romantic partner. Decision points such as these are pervasive and often highly consequential for one’s relationships with others. Prior work has shown that individuals have specific, stable preferences that guide their choices regarding familiar others (Guassi Moreira et al., 2020). Given that no two people we know are exactly alike, one might expect subjective representations of familiar others (i.e., one’s mental model of someone) play a key role in shaping social decision behavior. However, surprisingly little work has examined this. In the current study, we used functional magnetic resonance imaging (fMRI) and natural language processing (NLP) to examine the link between mental representations of familiar others and social decision behavior towards those individuals.

By and large, most social decision-making studies have not examined how mental representations of others shape behavior. Instead, studies typically investigate how cognitive or affective heuristics are applied over characteristics of choice alternatives to guide decision behavior. These are heuristics that help process a variety of decision-level features that convey information about risk, mood, ambiguity, inequity, harm, others’ perspectives, and so on (Austerweil et al., 2016; Cole & Bruno Teboul, 2004; Crockett et al., 2017; FeldmanHall & Chang, 2018; Hackel et al., 2017; Kao et al., 2023; Sampaio et al., 2023; Yu et al., 2019). Because so much attention has been devoted to such heuristics and their role in parsing decision-level features of choice alternatives, studies tend to disproportionately focus on contexts in which one’s choices impact strangers, anonymous others, and social agents with experimentally manipulated roles (Huettel & Kranton, 2012), despite the fact that everyday social decisions frequently implicate familiar others. Thus, the relationship between our mental representations of other people—especially known, familiar others—and the decisions we make regarding them remains largely unexplored.

Available evidence suggests that mental representations of other people should play a role in the decisions we make regarding those people. Several studies have delineated the neural architecture for thinking about other people (Broom et al., 2021; Hassabis et al., 2014; Mitchell et al., 2005; Yin Wang et al., 2017), and recent work has further extended this by investigating how distributed neural response patterns encode the identities and attributes of specific others (Chavez & Wagner, 2020; Hassabis et al., 2014; Yin Wang et al., 2017). This encoding constitutes the basis of mental representations of familiar others – i.e., models that allow us to simulate, forecast, and extrapolate the opinions, thoughts, feelings, and actions of others. Mental representations of others are thought to be evoked when predicting an individual’s thoughts or behaviors after having learned about their personality traits (Fleming & Slank, 2015; Hassabis et al., 2014; Mitchell et al., 2006), witnessing demonstrations of reward or punishment(Ho et al., 2021; Yu et al., 2019), and adherence to norms and conventions (Hawkins et al., 2021; Sarathy et al., 2017), all suggesting that mental representations of other people ought to at least partly drive social decisions. Despite this work, however, it remains unclear how such mental representations influence social choice preferences involving familiar others.

Here we probed if and how mental representations of familiar, close others are associated with behavior towards those people. We deliberately chose to investigate mental representations of familiar *close* others for several reasons. First, close others are most commonly implicated in social decision scenarios. Results obtained by investigating close others are more applicable to everyday human life – more so than a confederate, fictive individual, or celebrity. Second, relationships with close others can provide much richer representational content to map onto behavior. Third, examining choice preferences among individuals *within* the category of close others helps map behavioral tendencies in a finer-grained manner (as opposed to coarsely examining choice preferences between close versus non-close others) that is likely to capture the kinds of distinctions that pervade everyday life, where people must often consider the consequences of their choices for multiple familiar others (rather than deciding, for example, whether to favor a friend or a stranger). We specifically studied choice preferences between a parent and close friend because group-level choice preferences between parents and friends have been well documented(Guassi Moreira et al., 2018, 2020, 2021), and because these two particular relationship categories are motivationally salient across individuals and for much of the lifespan (Crone & Fuligni, 2020; Crosnoe & Johnson, 2011; Fuligni, 2019; Greenberg et al., 1983; Syed & Mitchell, 2013).

We assessed mental representations using multiple modalities of measurement by leveraging neuroimaging (specifically, fMRI) and applying NLP techniques to written text data. With fMRI, we based our analyses on key theories in social neuroscience that stipulate that one’s representation of the self serves as the template for all other mental models, and that self-relevance (i.e., high self-other representational overlap) is a key driver of behavior (Mitchell et al., 2006; Tamir & Mitchell, 2013). Thus, we examined the degree to which self-other representational overlap differed between parent and friends in brain regions related to social cognition and value-based processes—key psychological processes implicated in social decision-making—and then tested whether this self-other representational overlap tracked with social choice preferences. In particular, following extensive evidence from the established literature, the bilateral temporoparietal junction (TPJ) and dorsomedial prefrontal cortex (dmPFC) were probed as regions sensitive to social cognitive processing (Pfeifer & Peake, 2012; Saxe & Powell, 2006) and the medial prefrontal cortex (mPFC) and bilateral nucleus accumbens (NAcc) were specified as value-based processing regions (Delgado et al., 2016).

Using the linguistic data, we employed machine learning to test whether we could engineer a linguistic signature of social decision preferences from written text about one’s parent and friend (Fatima et al., 2021), essentially decoding choice preferences from one’s representation of others that is available in written text data. Finally, we combined the neuroimaging and linguistic data to test whether high correlation between representational similarities in the brain and language were associated with stronger choice preferences. We did so for two related reasons. First, information from one modality can help disambiguate findings from another. Second, generally, information captured with different methods may capture different aspects of one’s mental representations of other people, and using multiple modalities to measure mental representations of others can confer greater sensitivity to links between such representations and behavior.

A schematic overview of the entire study’s workflow is depicted in Figure 1. We first used neuroimaging data to test for differences in how the brain constructs representations for one’s parent and friend using the self as a reference (i.e., are friends or parents represented more similarly to the self?). We anchored representations in relation to the self because of aforementioned prior work in psychology and neuroscience showing that humans often use mental representations of the self as the basis for representations of others (Cadinu & Rothbart, 1996; Krienen et al., 2010; Mitchell et al., 2006; Todd & Tamir, 2024), and that perceived similarity to oneself is often linked to preferences regarding others (Fischer & Savranevski, 2023; Froehlich et al., 2021; Hackel et al., 2017). We then examined whether individual differences in the degree of representational similarity between oneself and particular close others were statistically associated with individuals’ social decision preferences between parents and friends. Next, we pivoted to linguistic data to examine whether social decision preferences regarding close others could be predicted from written descriptions of them. To do so, we validated a linguistic signature of choice preferences across several datasets with different types of social decision tasks and with additional characterizations of relationships (e.g., measures of relationship quality). Afterwards, we tested for high correspondence between information from both modalities (neural, linguistic) by correlating neural representations with linguistic signature scores. Finally, we finished by probing whether high correspondence between the two types of representations was related to stronger social decision preferences for a given other (i.e., if neural and linguistic representations both show a strong parent ‘bias’, is one even more likely to evince a parent-over-friend social choice preference?). In all, this study aims to reveal how mental representations of specific, close others influence social decision behavior towards those individuals.

**Figure 1.**
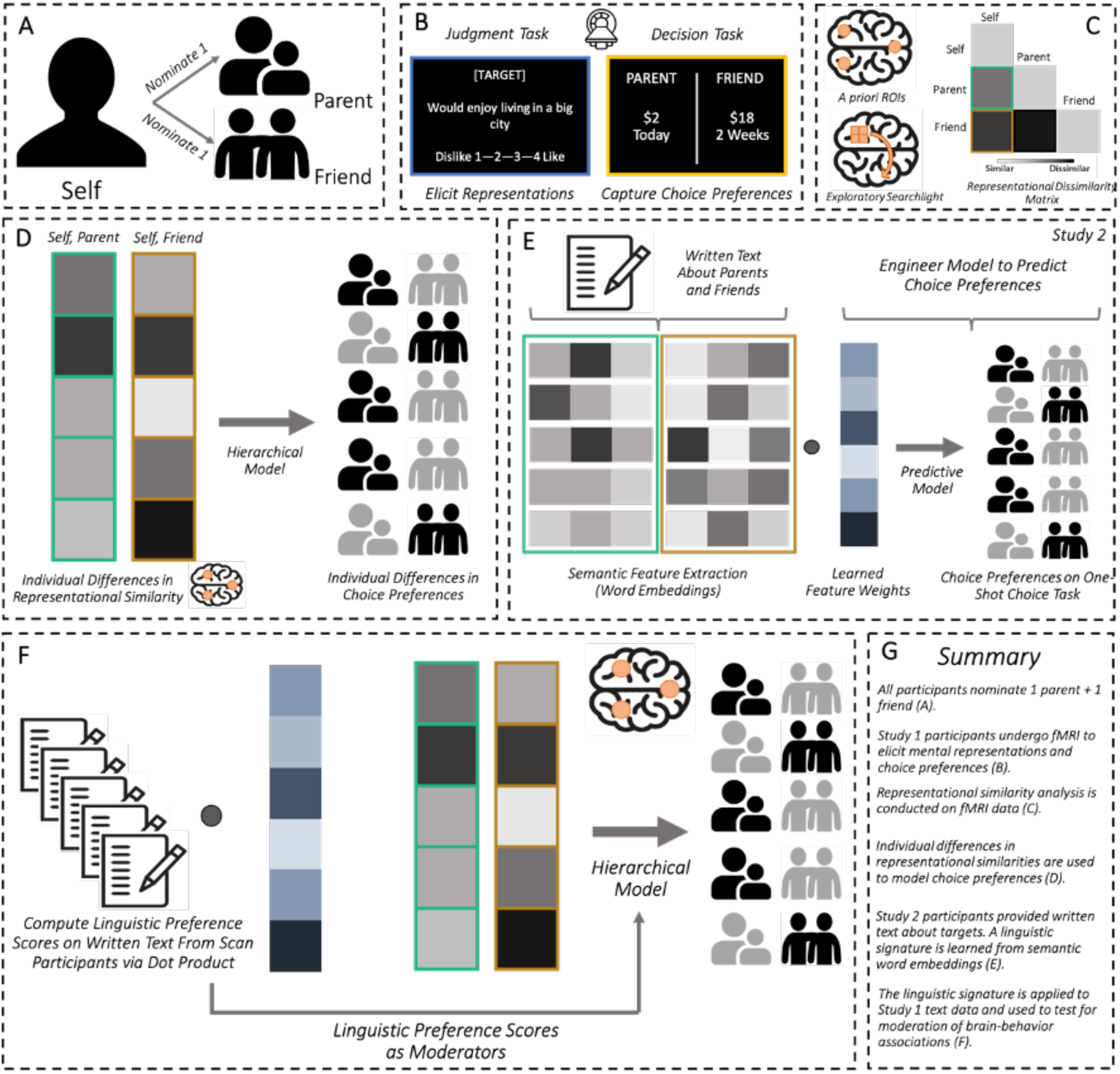
Study Overview. *Note.* (*A*) Participants in both studies nominated a parent and same-age friend of their choice who was not a current romantic partner or family member. (*B*) Participants in Study 1 underwent fMRI scanning while completing a task designed to elicit mental representations of the self, parent, and friend, as well as a social decision-making task that involved allocating monetary and social rewards between parent and friend. *(C)* RSA was used to compare parent and friend representations (directly, and relative to the self) in six *a priori* ROIs and in exploratory searchlight analyses. ROI mask visualizations can be accessed in Figure 1-1 of the Extended Data. *(D)* Hierarchical modeling was used to test for associations between individual differences in self - other (parent and friend) representational similarity in the ROIs and decision preferences between parents and friends. *(E)* Study 2 participants provided written memories about, and/or adjectives describing their nominated parent and friend and completed a one-shot social decision task. Semantic features for parent and friend text were extracted (Word2Vec embeddings) and used to create and validate a linguistic signature of participants’ decision preferences between parents and friends on the decision task. The linguistic signature yielded ‘linguistic preference scores’ that quantified implicit social choice preferences embedding in linguistic representations. These scores generalized to additional data (other social decision tasks, questionnaire-based measures of relationship characteristics, such as quality and social loss aversion). *(F*) The linguistic signature was used to compute linguistic preference scores on written text data obtained from Study 1 participants. Scores were entered as moderators in models predicting decision preferences from neural representational similarity. The study workflow is summarized in (*G*).

## Materials & Methods

### Study 1

#### Participants

A total of 63 (Mean age = 18.81 years, SD = 1.05 years, 73% female) individuals participated in the current study. Participants were recruited from the University of California, Los Angeles (UCLA) student body by tapping existing internal recruitment pools, classroom advertisements, and an email from the registrar’s office. Our *a priori* sample size goal was to recruit and run as many participants as possible during the winter and spring academic quarters (January – June, 2023). In order to be eligible for the study, participants must have been between 18-22 years at the time of enrollment and exhibit no MRI contraindications (e.g., claustrophobia, non-MR safe metallic implants, etc.). All participants were fluent in English. Four participants did not have imaging data available (3 due to software issues preventing data collection/reconstruction and 1 due to participant failure to fully disclose permanent jewelry during safety screening), rendering a sample size of *N* = 59 for imaging analyses. All participants provided consent in accordance with the policies of the UCLA Institutional Review Board. Participants were compensated $40 (USD) upon completion of the study. All data and materials are publicly available on the Open Science Framework (OSF; osf.io/fzds6).

Demographically, the full sample was comprised of 46 individuals assigned the female sex at birth (73%). In terms of gender, 43 individuals identified as women (68.3%), 17 as men (27.0%), 2 as non-binary (3.2%), and 1 (1.6%) declined to respond. Ethnically, 12 identified as Hispanic/Latinx (19.0%). Racially, 24 identified as Asian/American (38.1%), 23 identified as Caucasian (36.5%), 7 identified as mixed race/multi-ethnic (11.1%), 5 identified as another category (‘Other’, 7.9%), 1 declined to respond (1.6%), and 0 identified as Native Hawaiian/Pacific Islander and American Indian/Alaska Native (0%). In terms of sexual orientation, 46 individuals identified as straight/heterosexual (73%), 8 identified as bisexual (12.7%), 4 identified as homosexual (gay/lesbian, 6.3%), 3 identified as another category (‘Other’, 4.8%), and 2 identified as asexual (3.2%). Finally, 57 participants reported relying on their parents for financial assistance (90.5%). The self-reported median household income among participants was $100,000 (range = $18,000 - $3,000,000).

#### Protocol Overview

Participants completed the study in a one-hour session at UCLA’s Staglin One Mind Center for Cognitive Neuroscience. Participants were greeted by an experimenter upon arrival, consented, nominated a parent and close friend, received training for the two fMRI tasks, underwent task-based fMRI scanning, and then completed a post-scan survey. Major elements of this procedure are described in greater detail below.

#### Parent-Close Friend Nomination

Prior to receiving training for the fMRI tasks, participants were informed the purpose of the study was to understand how the brain encodes information about close others and how it uses such information when making choices about others. Participants were then asked to nominate a parent and their closest friend. The closest friend was not allowed to be a current romantic partner or a family member. For the parent, the majority of participants chose a mother (84%) whereas the remainder chose a father (mean age of nominated parent: 51.4 years, SD = 5.6). The majority (69%) of participants nominated a close friend assigned the female sex at birth (mean age of nominated parent: 19.1 years, SD = 1.2 years).

#### fMRI Acquisition Parameters

fMRI data were collected using a research-dedicated 3T Siemens Prisma scanner and a 32-channel head coil. Functional scan sequences were designed to be consistent with the sequences used by the Human Connectome Project (humanconnectome.org), which uses the University of Minnesota’s Center for Magnetic Resonance Research (CMRR) multi-band accelerated EPI pulse sequences (github.com/CMRR-C2P/MB). A high resolution T1 magnetization-prepared rapid-acquisition gradient echo (MPRAGE) structural image was acquired for registration purposes (TR = 1900 ms, TE = 2.48 ms, Flip Angle = 9°, FOV = 256 mm^2^, 1 mm^3^ isotropic voxels, 208 slices, A >> P phase encoding). Functional runs were comprised of T2*-weighted multiband echoplanar images (TR = 1240 ms, TE = 36 ms, Flip Angle = 78°, FOV = 208 mm^2^, 2 mm^3^ isotropic voxels, 60 slices, A >> P phase encoding, multiband acceleration factor = 4). Single-band reference images were acquired for each functional run.

#### Behaviors and Preferences Judgment Task

Mental representations of the participant (self), close friend, and parent were elicited during fMRI scanning with a computerized behaviors and preferences judgment task (Tamir & Mitchell, 2013; Thornton & Mitchell, 2018). The task entails rating how well a series of preferences describe a given target. On each trial, a label for a given target was displayed at top of the screen (‘SELF’, ‘PARENT’, or ‘FRIEND’), a preference was displayed in the center of the screen, and a 4-point Likert scale was displayed at the bottom (1 = “Strongly disagree”, 4 = “Strongly agree”). Examples of preferences include “really enjoys living in big cities”, “thinks modern art is uninteresting”, and “feels comfortable talking about personal issues”. Participants used a button box to indicate their response. Each trial was presented for 4500 ms, and was followed by a jittered fixation screen consisting of a black ‘+’ in the center of the screen (mean = ∼2000 ms, possible range = [200–10165 ms], distributed exponentially). Trials were ‘chunked’ by target such that participants would rate at least seven consecutive trials about a given target before moving on to another target to help reduce the cognitive load of rapidly switching between targets. To help ensure participants did not miss a transition between chunks of targets, the fixation cue at the center of the screen was displayed in red prior to the start of a new chunk.

Participants completed 225 trials total (75/target), spread across four runs. The number of trials per target varied across runs. Importantly, the exact same set of stimulus items (preferences) were used for each target (self, parent, friend), helping ensure that representations of oneself, a parent, and friend were comparably elicited and that different patterns of effects between self-parent and self-friend similarities would not be driven by stimulus differences. Each run typically lasted between 345–375 s, given the random jitter.

Preferences for this task were drawn from an open, online pool used in prior studies (https://jasonmitchell.fas.harvard.edu/links/; accessible on our OSF repository at osf.io/sfc6b). Using methods consistent with prior studies(Tamir et al., 2016), we narrowed the initial 800+ item pool into a final set of 75 items based on how strongly they were associated with Big Five personality traits (De Raad, 2000; John & Srivastava, 1999). We chose to use the Big Five as an organizing framework to structure our measurement of participant representations because personality traits are a key component of individual identities, are known to be broadly generalizable, and encompass other key types of information (e.g., they imply information about mental state-scenario associations). The optimization procedure involved applying a genetic algorithm to stimulus item ratings solicited in an independent, crowd-sourced sample. Further stimulus optimization details are hosted on our OSF page (osf.io/zcu7n). We checked whether reaction time data from the task differed between condition pairs of interest (self, parent; self, friend) while accounting for random effects of subject and stimulus by using hierarchical modeling. No such differences emerged. The task was programmed in jsPsych (de Leeuw et al., 2023).

#### Social Decision-Making Task

We used a pair of computerized delay discounting tasks to assess social decision preferences between parents and close friends. Our rationale for choosing a discounting task is threefold. First, discounting decisions are both pervasive in everyday life and are thought to be important for shaping life adjustment outcomes (Golsteyn et al., 2014; Lebeau et al., 2016). Second, computerized discounting tasks deployed in research settings are flexible in their configuration, allowing researchers to study social decision behavior in the face of different reward outcomes (Seaman et al., 2016). Last, we have previously shown that discounting tasks with social choice manipulations reliably tap decision preferences between specific social partners and are related to socioemotional facets of interpersonal relationships such as quality and subjective relationship value; we also previously observed that choice preferences from similar tasks tend to be correlated (Guassi Moreira et al., 2021).

Participants completed two separate runs of a delay discounting task. On each trial, participants were presented with two hypothetical scenarios that pitted outcomes for their nominated parent and friend against each other. One scenario involved a relatively immediate, smaller reward and the other involved a relatively delayed, larger reward. The delays could take the value of zero (‘Today’), 2 weeks, 4 weeks, and 6 weeks. Values of zero and 6 weeks were never presented in the delayed or immediate scenarios, respectively. The two runs differed in the type of rewards offered to participants. One of the runs offered participants hypothetical monetary rewards (in USD) to be earned on behalf of either nominee (e.g., $16 for a given target); the other run was comprised of social rewards, offering participants hypothetical time spent with either target (e.g., 16 minutes of time spent with a given close other). Participants were instructed to think of the latter condition as *additional* time they could spend with their parent or friend, combined with the time they already spend with either close other.

Numeric outcome values for each type of reward ranged from 2 – 30. There were 49 unique combinations of numeric values and time pairings (e.g., $2 now versus $18 two weeks from now) for each condition, resulting in 294 unique possible trial types (Guassi Moreira et al., 2021). In the interest of reducing in-scan task demands for participants, each run was comprised of 30 trial types randomly selected from the total 294. It was heavily stressed that participants were to complete the task as if rewards were real, even though they were hypothetical. Participants had 4500 ms to indicate their decision via button box. The trial terminated once they completed their response. Each trial was interspersed with a fixation stimulus ‘+’ and a random jitter (similar parameters as the previous task). Participants completed two runs of the task (1 monetary, 1 social). Each run lasted approximately 90–135 s, with variability owing to the self-paced nature of the task. The visual configuration of task stimuli were generally consistent with prior implementations (Guassi Moreira et al., 2021; Seaman et al., 2016). Both runs were programmed in PsychoPy (Peirce et al., 2019). This task was always presented to participants after the behaviors and preferences judgment task. Outcome types for this task were counterbalanced between participants.

#### Post-Scan Survey

Participants completed a post-scan survey immediately following the scan. During the survey, they provided basic demographic information (sex and age) about their nominated parent and friend, were asked to list approximately 10 adjectives or short phrases describing each close other, and answered four written prompts asking them to recall specific memories with each close other. The prompts asked participants to write about their (i) favorite memory with each close other, (ii) a time when their close other made them feel supported, (iii) a time when their close other let them down, and (iv) and a time when they let their close other down. Participants were asked to provide 3-5 sentences per each memory with as much detail as possible. Text about parents and friends were collected in separate blocks. Block orders were randomized between participants. The survey also contained measures of their relationship quality with both close others (see Study 2 methods), questions about how often they see each close other (actual and ideal), a measure of social loss aversion towards each close other (i.e., how upset one would be if they could no longer spend time with a given social partner (Guassi Moreira & Parkinson, 2024)), and two one-shot social decision-making items (a dictator game and a forced choice question about whom they would rather spend time with).

More information on these latter measures is fully disclosed on our OSF page (osf.io/zcu7n). Participants finally finished the survey by answering demographic items about themselves.

### Preprocessing and Statistical Analysis

#### MRI data preprocessing

*fMRIPrep* (version 22.0.2)—an open source, freely available tool based on functional neuroimaging software *Nipype* (Gorgolewski et al., 2011)—was used to perform preprocessing MRI preprocessing. The T1-weighted (T1w) image was corrected for intensity non-uniformity (INU) with N4BiasFieldCorrection (Tustison et al. 2010), distributed with ANTs 2.3.3 (Avants et al., 2008) (RRID:SCR_004757), and used as T1w-reference throughout the workflow. The T1w-reference was then skull-stripped with a *Nipype* implementation of the antsBrainExtraction.sh workflow (from ANTs), using OASIS30ANTs as target template. Brain tissue segmentation of cerebrospinal fluid (CSF), white-matter (WM) and gray-matter (GM) was performed on the brain-extracted T1w using fast (FSL 6.0.5.1) (Zhang et al., 2001). Brain surfaces were reconstructed using recon-all (FreeSurfer 7.2.0) (Dale et al., 1999), and the brain mask estimated previously was refined with a custom variation of the method to reconcile ANTs-derived and FreeSurfer-derived segmentations of the cortical gray-matter of Mindboggle (Klein et al., 2017). Volume-based spatial normalization to one standard space (MNI152NLin2009cAsym) was performed through nonlinear registration with antsRegistration (ANTs 2.3.3), using brain-extracted versions of both the T1w reference and the T1w template.

The following template was selected for spatial normalization: *ICBM 152 Nonlinear Asymmetrical template version 2009c* (TemplateFlow ID: MNI152NLin2009cAsym) (Fonov et al., 2009).

For each of the BOLD runs found per subject, the following preprocessing was performed. First, a single-band reference was used as a reference volume. Head-motion parameters with respect to the BOLD reference (transformation matrices, and six corresponding rotation and translation parameters) were estimated before any spatiotemporal filtering using mcflirt (FSL 6.0.5.1) (Jenkinson et al., 2002). The BOLD time-series were resampled onto their original, native space by applying the transforms to correct for head-motion. These resampled BOLD time-series will be referred to as *preprocessed BOLD in original space*, or just *preprocessed BOLD*. The BOLD reference was then co-registered to the T1w reference using bbregister (FreeSurfer) which implements boundary-based registration (Greve & Fischl, 2009). Co-registration was configured with six degrees of freedom. First, a reference volume and its skull-stripped version were generated using a custom methodology of *fMRIPrep*. Several confounding time-series were calculated based on the *preprocessed BOLD*: framewise displacement (FD), DVARS and three region-wise global signals. FD was computed using two formulations following Power (absolute sum of relative motions) (Power et al., 2014) and Jenkinson (relative root mean square displacement between affines)(Jenkinson et al., 2002). FD and DVARS are calculated for each functional run, both using their implementations in *Nipype* (following the definitions by Power et al. 2014). The three global signals are extracted within the CSF, the WM, and the whole-brain masks. Additionally, a set of physiological regressors were extracted to allow for component-based noise correction (*CompCor*) (Behzadi et al., 2007).

Principal components were estimated after high-pass filtering the preprocessed BOLD time-series (using a discrete cosine filter with 128s cut-off) for the two *CompCor* variants: temporal (tCompCor) and anatomical (aCompCor). tCompCor components were then calculated from the top 2% variable voxels within the brain mask. For aCompCor, three probabilistic masks (CSF, WM and combined CSF+WM) are generated in anatomical space. The implementation differs from that of (Behzadi et al., 2007). in that instead of eroding the masks by 2 pixels on BOLD space, a mask of pixels that likely contain a volume fraction of GM is subtracted from the aCompCor masks. This mask was obtained by dilating a GM mask extracted from the FreeSurfer’s *aseg* segmentation, and it ensures components are not extracted from voxels containing a minimal fraction of GM. Finally, these masks are resampled into BOLD space and binarized by thresholding at 0.99 (as in the original implementation). Components are also calculated separately within the WM and CSF masks. For each CompCor decomposition, the *k* components with the largest singular values are retained, such that the retained components’ time series are sufficient to explain 50 percent of variance across the nuisance mask (CSF, WM, combined, or temporal). The remaining components were dropped from consideration. The head-motion estimates calculated in the correction step were also placed within the corresponding confounds file. The confound time series derived from head motion estimates and global signals were expanded with the inclusion of temporal derivatives and quadratic terms for each (Satterthwaite et al., 2013). Frames that exceeded a threshold of 0.5 mm FD or 1.5 standardized DVARS were annotated as motion outliers. Additional nuisance timeseries were calculated by means of principal components analysis of the signal found within a thin band (crown) of voxels around the edge of the brain, as proposed by (Patriat et al., 2017). The BOLD time-series were resampled into standard space, generating a *preprocessed BOLD run in MNI152NLin2009cAsym space*. First, a reference volume and its skull-stripped version were generated using a custom methodology of *fMRIPrep*. All resamplings can be performed with a single interpolation step by composing all the pertinent transformations (i.e. head-motion transform matrices, susceptibility distortion correction when available, and co-registrations to anatomical and output spaces). Gridded (volumetric) resamplings were performed using antsApplyTransforms (ANTs), configured with Lanczos interpolation to minimize the smoothing effects of other kernels. Non-gridded (surface) resamplings were performed using mri_vol2surf (FreeSurfer).

#### Multivariate Pattern Estimation

Multivariate patterns were estimated by modeling imaging data from the preferences task using a General Linear Model (GLM). Each run of the task was modeled using a fixed effects GLM in FSL. Every GLM consisted of the following explanatory variables: A regressor for each of the self, parent, and friend trials where participants rated the extent to which a given item described said target. The duration of each regressor corresponded to the length of the trial (4500 ms). These regressors were convolved with the hemodynamic response function (double gamma). The temporal derivatives of each were added to account for slice timing effects. The following confound regressors were included to statistically control for noise and head motion. Timeseries from the CSF+WM mask automatically extracted by *fMRIPrep* was added into the model, in addition to the tCompCor component. 24 head-motion regressors and individually censored volumes exceeding the FD and DVARS thresholds, both computed by *fMRIPrep*, were also included. Three linear contrasts were computed: [self - baseline], [parent - baseline], and [friend - baseline].

#### Representational Similarity Analysis

We used representational similarity analysis (Kriegeskorte et al., 2008) to summarize neural representations of others. RSA is useful for determining how various forms of psychological or behavioral information are encoded in the brain, and broadly, for comparing representational structure across modalities. Here we use RSA to examine how information about parents and friends is encoded in the brain (specifically self, other overlap), as well as how this information is relevant for social decision-making. The former set of analyses involved conducting various searchlight and ROI analyses; the latter used individual RDM cells to estimate statistical relationships between individual differences in neural self, other overlap and social decision preferences. These analyses are described in the following three sections.

#### A Priori ROI Analyses

We implemented RSA on region-of-interest (ROI) data to determine whether there was a ‘bias’ in the encoding of parent or friend representations in six brain regions (mPFC, dmPFC, L NAcc, R NAcc, L TPJ, R TPJ), defined *a priori* via meta-analytic maps for cortical regions and probabilistic atlases for subcortical regions (NAcc).

Cortical regions were identified by their peak meta-analytic coordinates. We did this by constructing an RDM based on between-run distances in multivariate patterns of brain activity. Between-run distances were used to avoid the potential confound of temporal “pattern drift”, which has been shown to persist even after voxel-wise detrending has been performed, but which can be alleviated by focusing on comparisons between patterns collected in different runs (Alink et al., 2015). Focusing on between-run multivoxel pattern comparisons is also recommended in designs such as that used in the current study, where particular trials (e.g., those corresponding to a particular target) are “chunked” together (Mumford et al., 2014).

Concretely, this meant that multivoxel patterns for self, parent, and friend were extracted for a given ROI (using t-statistic images) and distances were computed between patterns from each run for all possible target pairings. Because we were using between run distances, this also meant comparing between run patterns for the same target (e.g., the self condition for run 1 was compare to same condition in run 2). We ran analyses with both (1 – Pearson’s *r*) and Euclidean distances as the RDM-defining distance metric to check for robustness—analyses were consistent between the two metrics. Euclidean distances are reported in the main document.

The mPFC, dmPFC, and TPJ ROIs were defined by locating the respective brain regions on the Neurosynth meta-analytic platform (Yarkoni et al., 2011), noting coordinates for the global maximum (or within each hemisphere for the TPJ), and drawing a 5 mm radius sphere around the coordinates. The NAcc masks were defined using the Harvard-Oxford subcortical atlas as available in FSL. ROI masks are visualized in the Extended Data (Figure 1-1). We then computed pairwise differences between the [self and parent] and [self and friend] RDM cells for each of the six ROIs. Paired differences were modeled as being drawn from a Gaussian distribution such that *Diff_i_* ∼ *N*(δ*σ, σ^2^). The mean was parameterized as δ*σ so that draws from the distribution are in “native” units but summary statistics of paired differences are in the Cohen’s *d* metric (i.e., mean/standard deviation). Prior distributions were assigned to both δ 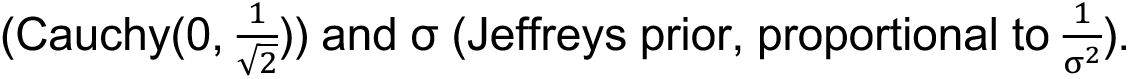 The Bayesian modeling software *stan* (Stan Development Team, 2020) was used to run these analyses via the *rstan* R package (no thinning, 4 chains, 2,000 samples/chain, 1,000 discarded burn-in samples).

#### Exploratory Searchlight Analyses

We conducted a set of exploratory searchlight analyses (Kriegeskorte et al., 2006) to identify which brain regions showed the greatest differentiation among the three representations (self, parent, friend) in a data-driven manner. The searchlight was meant to complement the ROI approach by ensuring we did not miss any additional brain regions that differentiated between targets. A 125 mm^3^ sphere (2 mm radius, excluding center voxel [(2 * 2) + 1]^3^) was moved throughout each participant’s brain and multivoxel patterns to each target were compared at each searchlight center. The first comparison we performed was a direct comparison between parent and friend representations. This was accomplished by extracting multivariate patterns for the parent and friend *t*-statistics at each center voxel, vectorizing the voxels within the sphere, and taking the (Pearson) correlation distance (1 – *r*) between the two patterns. As before, we only used between-run comparisons.

We used between-run distances within-target as a baseline. All between-run distances were placed into a vector and standardized. The mean of within-target distances (e.g., (parent run 1, parent run 2), (friend run 2, friend run 4)) was then subtracted from the mean of (parent, friend) distances. We conducted a second series of analyses comparing representational overlap between the self and each target. Specifically, distances between neural representations of one’s self and one’s parent were compared to distances between representations of one’s self and one’s friend in a procedure similar to that described above.

#### Modeling Social Decision Preferences

We used Bayesian statistics for all regression models involving choice behavior on the social decision-making tasks. Given that there exists notable intra-and inter-individual variability in decision-making behavior, we chose to model the data at the trial-level, necessitating the use of a hierarchical logistic regression model. While a hierarchical model is not strictly needed to handle nested data (McNeish et al., 2017), we use it here because it allows us to flexibly model both within-and between-participant sources of variance while regularizing the model coefficients to enhance out-of-sample generalizability. The form of the within-person component of the model is notated below.

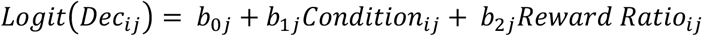

This is the core component of the model because it encodes social decision preferences between parents and friends. Here, the *i*-th decision (*Dec*, 1 = discount, 0 = non-discount) from the *j*-th participant was modeled as a function of a *Condition* dummy code and a *Reward Ratio* variable. *Condition* is coded such that 0 = friend outcome associated with discounting option, parent outcome associated with non-discounting option and 1 = parent outcome associated with discounting option, friend outcome associated with non-discounting option. The coefficient associated with *Condition* (*b_1j_*) encodes social decision preferences between parents and friends because it reflects the mean difference in choice tendencies between the two conditions (after statistically adjusting for other terms in the model). A *b_1j_* coefficient > 0 indicates a preference towards parents: there is greater likelihood of choosing to discount when the parent outcome is associated with the smaller immediate reward, *and* a greater likelihood of not discounting when parents are affected by the larger delayed reward. A *b_1j_* coefficient < 0 indicates a friend-oriented preference following the same logic: a reduced likelihood of choosing to discount when the friend outcome is associated with a larger delayed reward *and* an increased tendency of discounting when associated with the smaller immediate reward. *Reward Ratio* is the quotient of the non-discounted outcome over the discounted outcome and is a key covariate for several reasons. Primarily, it serves as a ‘yardstick’ to gauge whether representational information is meaningfully related to decision preferences beyond trial-level features. This is notable because various cognitive heuristics used in decision-making are typically applied over such features; adjusting for this here helps ensure that representational information is still related to choice behavior over and above other cognitive processes. In addition, controlling for *Reward Ratio* helps to make choices more comparable between trials, and provides a ‘sanity check’ for participant engagement (greater reward ratio should generally be associated with a reduced likelihood of discounting regardless of condition). Additionally, a *post hoc* examination revealed that trial-level task features (e.g., reward ratio, delay ratio, magnitude of rewards) were not significantly oversampled in either choice condition, ruling out the possibility that social decision preferences were not conflated with trial-level choice option features. All three terms in the within-person component of the model were allowed to vary randomly across participants.

Allowing the *b_1j_* coefficient to vary randomly across participants meant that we could model variability in individual decision preferences. We did so across various models by entering different between-participant variables. These between-participant variables are statistically allowed to interact with the Condition dummy code (i.e., a cross-level interaction in a one-level model), but are substantively conceptualized as the association between participant-level individual differences and social choice preferences. The following sets of between-person predictors were tested (each group in a separate model): (i) dissimilarity values for self and parent, as well as self and friend calculated from behavioral ratings, (ii-v) neural dissimilarities from each ROI (bilateral NAcc and TPJ ROIs were entered in the same model). Here we notate the between-participants portion of the model with neural dissimilarity values from mPFC as an illustrative example (interested readers can consult our publicly available code for further model details).

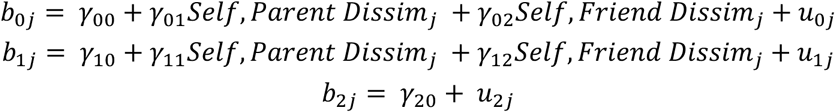

Analyses involving linguistic preference scores (described in the next section) were entered as part of a three-way interaction between neural dissimilarity and *Condition* in subsequent analyses. All continuous predictors were grand mean centered prior to running the model. All models were implemented using the *brms* package in the R statistical software library (no thinning, 8 chains, 2500 samples/chain, 1250 discarded burn-in samples). A weakly informative standard normal prior was placed on all slope coefficients and brms default priors were used for all other model parameters.

Because the output of Bayesian analyses is a distribution, we performed statistical inference in a graded manner and thus do not report nor use p-values to conduct frequentist inference. Instead, we evaluated the weight of statistical evidence using a combination the Region of Practical Equivalence (ROPE) method (Kruschke, 2011, 2013) and noninferiority testing (Spiegelhalter et al., 1994; Wiens, 2002). Traditionally, the ROPE method entails specifying a symmetric region in parameter space around the null value, calculating a credible interval (CI), and then evaluating whether the CI intersects at all with ROPE. CIs that do not overlap with ROPE result in a rejection of the null hypothesis, CIs that are entirely contained within ROPE result in acceptance of the null, and any other configuration falls in undecided.

Originating in clinical trials research, noninferiority testing is similar: if the CI partly overlaps with the upper limit of ROPE but does not pass through below the lower limit (regardless of its relation to the null value), the parameter value is declared to be *noninferior* to the null value (the same method can be used in the other direction, e.g., for ‘nonsuperior’ inference).

Here we combine these methods in the following manner. If the CI does not overlap with ROPE, we deem the strength of evidence to be robust. If the CI does not overlap with the null value (usually zero), but partly overlaps with ROPE, we deem the strength of evidence to be moderate. If the CI overlaps with ROPE and the null value (in the manner usually described in noninferiority testing), we deem the strength of evidence to be modest. If the CI completely *spans* ROPE (i.e., overlaps and exceeds both ends of ROPE), deem the evidence to be inconclusive. If the CI is contained within rope, we interpret the finding to support the null result. We deliberately follow these inferential criteria because it displaces the inferential focus from a problematic binary ‘significant vs not-significant’ label (Kruschke, 2018) to a more nuanced conclusion describing the *degree* of evidence in relation to meaningful benchmarks. We conducted a permutation analysis on several study variables to determine whether zero was an empirically reasonable null value to use. Analyses showed the expected value of a correlation coefficient under a permuted distribution was indeed zero. Posterior distributions were summarized with the mean of the relevant statistic (e.g., Cohen’s d, regression coefficient) and an 89% highest density credible interval (HDI; all samples within the interval have a higher density than those outside it). We used 89% credible intervals upon the recommendation that wider intervals (e.g., 95%) are more sensitive to Monte Carlo sampling error (Makowski et al., 2019; McElreath, 2015).

### Study 2

#### Datasets

Study 2 was comprised of secondary analysis of three large existing datasets (*N* = 1,641 unique participants total) containing written text about parents and friends (nominated using the same procedure as described in Study 1) as well as social decisions involving these two others. We used one dataset (Dataset 1) to engineer a linguistic signature of social decision preferences. The written text data and social decision measures in this dataset were collected as part of a broader study aiming to answer an orthogonal research question (Guassi Moreira & Parkinson, 2024). The other two datasets (Dataset 2, Dataset 3), and additional social decision measures from Dataset 1, were used to validate the linguistic signature by ensuring that it predicted social decision preferences in additional contexts. These datasets come from a series of studies on social decision-making conducted by the research team (Guassi Moreira et al., 2018, 2020, 2021). These prior studies have yielded published results, but the written text data have not been analyzed before. A detailed description of each dataset follows.

Dataset 1 was comprised of five independently collected subsamples (S1.1 – S1.5). Participants in each subsample were asked to nominate a parent and close friend of their choice and provide approximately ten adjective or short phrases (e.g., ‘happy go lucky’, ‘pain in the butt’) describing each close other. Participants in S1.1 were also prompted to recall four types of memories involving each social partner – instances in which they felt supported by the close other, in which they supported their close other, in which their close other let them down, and their favorite memory with their close other. Social decision preferences were assessed with a one-shot dictator game question in which individuals allocated a hypothetical $100 pot between parents and friends in $5 increments, and another in which they indicated who they would rather spend time with given a day with no other obligations or commitments. Subsamples S1.1 – S1.5 totaled *N* = 870 participants (range of subsample size *n* = 45 – 454). Subsamples were recruited from the UCLA undergraduate psychology subject pool and online crowdsourcing platforms (Mturk, Prolific) in 2022. All prior analyses involving the social decision-making data from this dataset were recently published (Guassi Moreira & Parkinson, 2024).

Dataset 2 was comprised of five independently collected subsamples (S2.1 – S2.5) as a part of a series of studies on social decision-making. Participants in each subsample were asked to nominate a parent and close friend of their choice, write a memory they share with each target, and list several words or phrases describing the target. They were then administered a social decision-making task that forced them to make choices between the parent and friend. Social decision preferences were assessed with a version of the Columbia Card Task (Figner et al., 2009)—a multi-trial risk-taking paradigm—that was modified to capture social decision preferences when making choices with conflicting outcomes between a pair of others (Guassi Moreira et al., 2018). Subsamples S2.1 – S2.5 totaled *N* = 574 participants (range of subsample size *n* = 46 – 225). Subsamples were recruited from the UCLA undergraduate psychology subject pool and the broader West Los Angeles community between 2016 - 2020.

Dataset 3 was comprised of four subsamples (S3.1 – S3.4). Some of the participants in these subsamples also provided data for subsamples in Datasets 1 and 2. The distinctive contribution of this dataset relative to the others is that it uses a different social decision-making paradigm. As with the other datasets, participants were prompted to nominate a parent and close friend. Some subsamples followed the text generation procedure described for Dataset 1, others follow the procedure for Dataset 2, meaning that all subsamples provided adjectives and short phrases describing each other, and some also provided a memory of each other. Social decision preferences were assessed with the same task as in Study 1: a multi-trial delay discounting task that was modified to capture social decision preferences when making choices with conflicting outcomes between a pair of others (Guassi Moreira et al., 2021). Subsamples S3.1 – S3.4 totaled *N* = 920 participants (range of subsample size *n* = 61 – 454). Subsamples were recruited from the UCLA undergraduate psychology subject pool (2018-2019) and from an online crowdsourcing platform (2022).

Further information about each dataset, such as participant demographics and methodological details, can be accessed in the original publications cited above. Self-reported relationship quality between participants and their nominated parents and friends was collected in all three datasets. All participants provided consent in accordance with the policies of the UCLA Institutional Review Board. Depending on their method of recruitment, participants were either compensated in course credit or monetary remuneration (USD). All study materials and code are publicly available on the Open Science Framework (OSF; osf.io/fzds6). For privacy reasons, we are unable to share the raw or preprocessed text data.

The following two sections detail (i) how we used Dataset 1 to engineer a linguistic signature of social decision preferences based on one social decision measure and then (ii) how we used data from all three datasets to further validate the signature to verify that scores derived from the signature were related to social decision preferences in novel data and novel social decision measures.

### Engineering a Linguistic Signature via Machine Learning

Since our goal was to examine whether linguistic representations of close others encode information about social decision preferences, we trained a model to predict social decisions from semantic features of written text data. We sought to produce a linguistic signature of social decision preferences that followed the logic of neural signatures and pattern expression analysis in the cognitive and affective neuroscience literature (Chang et al., 2015; Wager et al., 2013), both in the process used to engineer the signature and its application. The first step in this process was to test candidate models with cross-validation on the subsamples from Dataset 1 to settle on model specifications (type of model, hyperparameters). The pipeline for this task was comprised of the following steps: (i) social decision measure selection, (ii) text preprocessing, (iii) representation extraction, (iv) leave-one-out cross-validation using each subsample, and (v) estimating model weights on the entire dataset using the model with the best cross-validation accuracy.

### Selecting a Social Decision Measure for Signature Construction

We used the forced choice question about which close other the participant would rather spend a free day with (coded such that parent = 1, friend = 0) as our measure of social decision preferences. We chose this metric for assessing social decision preferences for four reasons. First, the binary nature of the item makes for an intuitive classification metric with which to evaluate model performance (percent accuracy). Second, using a binary forced-choice measure prevents participants from being impacted by a neutral scale response bias and requires them to disclose an unambiguous preference. Third, framing social decision preferences in terms of spending time (a finite resource subject to opportunity cost) with others simulates a real-world scenario that could generalize better to behaviors outside of the lab. Last, the simplicity of the item may help it better translate to other, more complex measures of social decision-making. Preferences for parents over friends for each sample follows: S1.1 = 51% chose parents over friends; S1.2 = 42%; S1.3 = 42%; S1.4 = 46%; S1.5 = 42%.

### Text Preprocessing

Written text data was preprocessed in a manner consistent with other similar studies (Fatima et al., 2021), implemented with the TextBlob python library. The following preprocessing pipeline was applied to each participant’s data. We first removed stopwords, as defined by TextBlob, in addition to the words ‘favorite’ and ‘memory’ (as many participant responses to memory prompts began with this phrase). Afterwards, each word was forced into its lowercase form and lemmatized. Words were then tagged as parts of speech and filtered. Only nouns, adjectives, verbs and adverbs were in the preprocessed word list, corresponding to the following TextBlob tags:’JJ’,’JJR’,’JJS’,’NN’,’NNS’,’NNP’,’NNPS’,’RB’,’RBR’,’RBS’,’VB’,’VBD’,’VBG’,’VBP’,’VBZ’. Parent and friend memories were preprocessed separately, meaning that each participant had a set of a preprocessed words from written text data about their nominated parent, and another preprocessed word set about their friend.

### Representation Extraction

We used word embeddings to extract linguistic representations of parents and friends from participants’ written text data. This approach quantified semantic meaning from the preprocessed word set in a bottom-up fashion using pretrained Word2Vec word embeddings. We extracted word embedding vectors for each word in the parent and friend word sets separately, and then averaged the vectors of all set words together. We used the pretrained embeddings from Google’s 300-dimension Word2Vec model (Mikolov et al., 2013), as implemented in the genism python package. Words in the word set that did not have a corresponding word embedding vector were excluded from the representation. In this approach, the semantic content of each participant’s text data for a given close other (parent, friend) was summarized as a composite 300-dimensional vector. We used principal components analysis (PCA) to reduce dimensionality in order to aid modeling. Both parent and friend composite vectors for each subject were reduced to 30 dimensions, consistent with similar work (Yingying Wang et al., 2023). On average, 30 dimensions accounted for approximately 83% of the variance in written text data, collapsing across parent and friend text in all datasets.

### Leave-One-Subsample-Out Cross-Validation

We used leave-one-subsample-out cross-validation to determine the best specifications for defining a linguistic signature of social decision preferences. We specifically sought to identify the best type of machine learning model and set of hyperparameters. We identified three possible types of machine learning models: a support vector classifier (SVC), a ridge regression classifier (RRC), and regularized logistic regression (RLR). Hyperparameters for each model were selected using a grid search within each iteration of cross-validation. The candidate values for all possible hyperparameters (C, γ, α) were {1, 5, 10, 50, 100}.

Our cross-validation approach took advantage of the independent subsamples in Dataset 1—we chose to leave-one-subsample out for each iteration of cross-validation. Since the goal of cross-validation is to estimate out-of-sample modeling performance, we reasoned the natural partitioning of each subsample as independent datasets best served this goal. This meant that each iteration of cross-validation proceeded such that one subsample was scaled and set aside, the other four were aggregated and scaled, a grid search was performed to find the best hyperparameter(s), the best hyperparameter was then used to estimate model weights on the four aggregated subsamples, and the model weights were finally applied to the left-out subsample to estimate the predictive accuracy of the model. This process was repeated five times, leaving one of the subsamples out each time, for all of the three models described above. This workflow was implemented using functions from the sci-kit learn python library.

### Estimating Model Weights

As previously noted, the binary forced choice question about spending time with either a parent (coded as 1) or friend (coded as 0) served as our criterion.

The principal components derived from both the parent and friend text data (60 total) were entered into the winning model from the prior step as predictors. The ensuing 60 coefficients (weights) from the model indicate how a ‘preference’ towards one close other is embedded in the linguistic representation and thus comprise the linguistic signature.

### Validating the Linguistic Signature

Once we engineered in our linguistic signature in Dataset 1, we set out to further validate it by correlating linguistic preference scores derived from the signature with other measures of social decision-making across the three datasets. The logic for doing so is that if our linguistic signature is truly capturing social decision preferences between parents and friends, then the signature should generalize to novel measures and predict preferences across different measures in the three datasets. To that end, we used the weights from our signature to compute preference scores and test whether those preference scores could predict social decision preferences on previously untested social decision tasks. Linguistic preference scores were calculated by taking the dot product between the model weights that comprised the linguistic signature and the feature-reduced text data from a given subject. The feature-reduced text data were the same as described above: PCA-reduced composite word2vec embeddings for parents and friends. When validating the linguistic signature, we used decisions from a one-shot dictator game where participants had to directly allocate money between parents and friend (Dataset 1), a multi-trial decision-making task that pitted gains and losses for parents against those for friends in a risky context (Dataset 2), and a multi-trial decision-making task where participants had to choose to allocate money for, or time spent with, their parents and friends in a delay discounting context (Dataset 3; the same task used in the Study 1). Finally, because social decision preferences are often predicted by attitudes towards social partners, we also tested whether the linguistic signature could predict relationship quality scores for parents and friends (available in all datasets). These validation tests for each outcome are described in greater detail below. Bayesian statistics were used for all analyses. The inferential criteria from Study 1 were applied to Study 2 analyses.

### Social Preferences in the One-Shot Dictator Game

The one-shot dictator game, collected in Dataset 1, asked participants to allocate a hypothetical pot of $100 between their nominated parent and friend. Values varied by increments of $5, ranging from [$100 parent, $0 friend] to [$0 parent, $100 friend], resulting in a 21-point ordinal variable. Responses were recoded to be the percentage of money allocated towards the parent (0.0, 0.05, 0.10… 0.90, 0.95, 1.00). A robust Pearson correlation between preference scores and dictator game allocations was computed using the Bayesian software package stan. We note that while the text data for this validation step is not novel (i.e., were used in model engineering), the outcome variable (dictator game allocations) is, and it is valuable to know whether preference scores can predict decision preferences in different contexts within-participants.

### Social Preferences in a Risky Context

Linguistic preference scores were next validated by determining whether they could predict parent vs friend preferences in a risky decision context. Participants in this dataset (Dataset 2) completed a multi-trial decision task that required them to make choices between a risky and a safe outcome. The risky outcome carried the potential for either a hypothetical monetary gain or a hypothetical monetary loss according to a known probability; the safe outcome was neutral (no gain or loss). The interests of parents and friends were pitted against each other such that any potential gain from a risky choice on a given trial would be credited to one partner, whereas any potential loss would be credited to the other partner. Participants completed multiple trials across two runs of the task, one in which risky gains benefitted one partner (e.g., parent) while the losses were incurred by the other partner (e.g., friend), and another run in which the opposite was true. Hierarchical Bayesian logistic regression was used to model the trial-level likelihood of making a risky (vs safe) decision as a function of a parameter that encoded choice preferences on the task and other trial-level covariates. We refer readers to (Guassi Moreira et al., 2018) for details about the form of the within-person component of the model. We entered preference scores derived from the linguistic signature as a between-person predictor to the model, allowing it to interact with the choice preference parameter from the model. Because the choice preference parameter was allowed to vary randomly across subjects, the ensuing interaction term codes for how choice preferences on the task vary as a function of preference scores derived from the linguistic signature. The brms package was used to fit the models (4 chains, 1000 discarded burn-in samples, 2000 samples/chain, no thinning).

### Social Preferences in a Delay Discounting Context

Linguistic preference scores were further validated by testing whether they could predict parent vs friend preferences in a delay discounting context. This is the same task used in Study 1, with the only difference being that more trials were administered to participants. Hierarchical Bayesian logistic regression was used to model the trial-level likelihood of making a discounting decision (i.e., choosing the immediate over delayed reward) as a function a parameter that encoded choice preferences on the task and another trial-level covariate (same within-person model as Study 1). We ran separate models for choices with monetary and social outcomes. Preference scores derived from the linguistic signature were entered into this model in the same manner described above. The brms package was also used to fit the models (using the same specifications that are listed above).

### Predicting Relationship Quality Using the Linguistic Signature

Last, we opted to relate preference scores derived from the linguistic signature to self-reported relationship quality with parents and friends. In doing so, our goal was to subject the linguistic signature to another validatory step that involved predicting *attitudes* that may also influence social decision behavior. A meaningful result for this test would indicate that motivationally relevant information about one’s attitude towards a close other is encoded in their linguistic representation.

Information about the self-report measure used here (Inventory of Parent and Peer Attachment) and its reliability is included in the Extended Data.

## Results

### Information about social decision preferences is embedded in neural representations of close others

*Different Categories of Close Others are Differentially Represented Across Brain Regions.* We compared the degree to which individuals construct mental representations of parents and friends in terms of self-relevance (i.e., representational overlap between self-parent compared to self-friend) in six *a priori* brain regions of interest (Table 1; Figure 2) that are involved in social processing and social decision-making (ROIs visualized in Figure 1-1). We found modest evidence to suggest greater self-other overlap for parents than for friends in bilaterally in the TPJ and NAcc. In contrast, there appeared to be greater self-other overlap for friends than for parents in mPFC.

**Figure 2.**
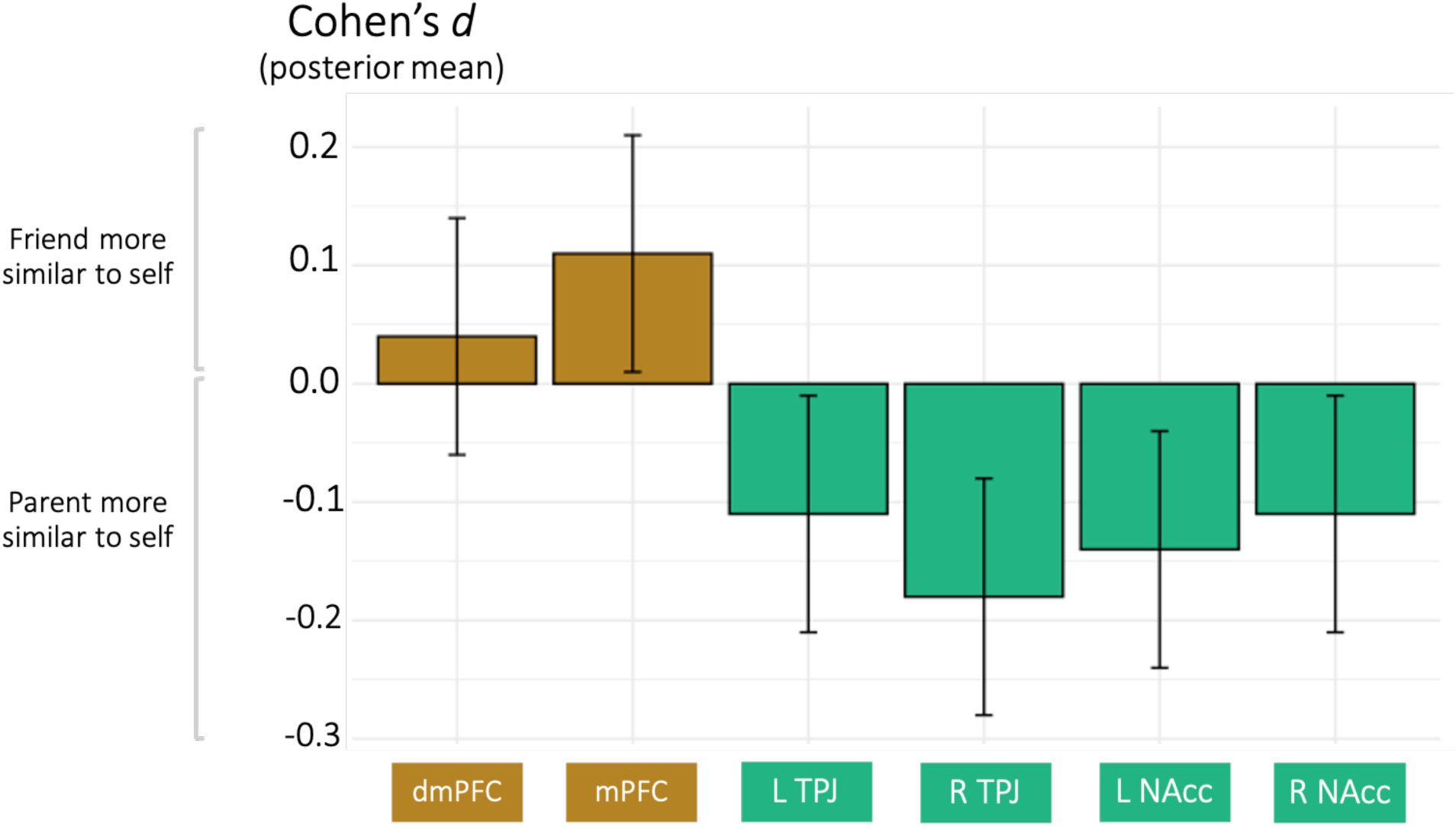
Brain regions involved in social cognition and value-based processes differentially construct representations of parents and friends in relation to the self. *Note.* ‘dmPFC’ refers to dorsomedial prefrontal cortex. ‘mPFC’ refers to medial prefrontal cortex. ‘TPJ’ refers to temporoparietal junction. ‘NAcc’ refers to nucleus accumbens. ‘L’ refers to left and ‘R’ refers to right. Cohen’s *d* values reflect paired differences between (self, parent) - (self, friend) dissimilarities of multivoxel response patterns extracted from the six brain regions displayed here. Error bars correspond to posterior standard deviations. Exploratory searchlight results to complement this analysis are depicted in Figures 2-1 and 2-2 in the Extended Data. Figure 2-3 of Extended Data shows the results of mass univariate analyses for comparison.

**Table 1.** Friend representations are more similar to self representations in cortical midline regions, whereas parent representations are more similar to self representations in the TPJ and NAcc.

| ROI | Paired Difference [89% HDI] |
| --- | --- |
| dmPFC | 0.04 [-0.15, 0.25] |
| mPFC | *0.11 [-0.10, 0.30]* |
| L TPJ | *-0.11 [-0.31, 0.09]* |
| R TPJ | *-0.18 [-0.38, 0.03]* |
| L NAcc | *-0.14 [-0.35, 0.08]* |
| R NAcc | *-0.11 [-0.30, 0.10]* |

*Note.* ‘dmPFC’ refers to dorsomedial prefrontal cortex. ‘mPFC’ refers to medial prefrontal cortex. ‘TPJ’ refers to temporoparietal junction. ‘NAcc’ refers to nucleus accumbens. ‘L’ and ‘R’ refer to left and right hemisphere, respectively. Values in the table reflect Cohen’s *d* statistics (posterior mean) for paired comparisons between dissimilarities of multivoxel representations of self and parent, and self and friend, in each brain region. Neural dissimilarities were calculated using Euclidean distance. Values in brackets reflect 89% highest density credible intervals drawn around the posterior distribution. ROPE was determined as [-0.1, 0.1]. Differences where the HDI partially overlaps with ROPE and includes 0 (indicating modest evidence for a difference between self-other overlap for friends vs parents) are italicized.

We ran two exploratory searchlight analyses to determine if brain regions other than the *a priori* ROIs also distinguish between familiar others. The first of these analyses tested for distinct representations of parents vs. friends and the second tested for differences in representational overlap between self and parent, and self and friend. Results of the first searchlight suggested that the precuneus contains distinct representations of close others (Figure 2-1). Exploratory examination of uncorrected results of this analysis revealed more extensive results in the precuneus and several smaller clusters in medial prefrontal cortex (mPFC; see Figure 2-2). The second searchlight yielded no significant results. Technical details of the searchlight are hosted on our OSF page (osf.io/zcu7n).

A traditional univariate analysis comparing the magnitude of brain activity evoked when answering items for each target ([self – parent], [self – friend], [parent – friend]) is also included in the Extended Data (Figure 2-3). This analysis revealed robust activity along the cortical midline and TPJ for both the [self – parent] and [self – friend] contrasts. For thoroughness, we analyzed behavior-based representations using the task’s Likert scale ratings and found greater self-other overlap for friends than for parents (posterior Cohen’s *d* = 0.50, [0.28, 0.71] 89% highest density credible intervals).

**Figure 3.**
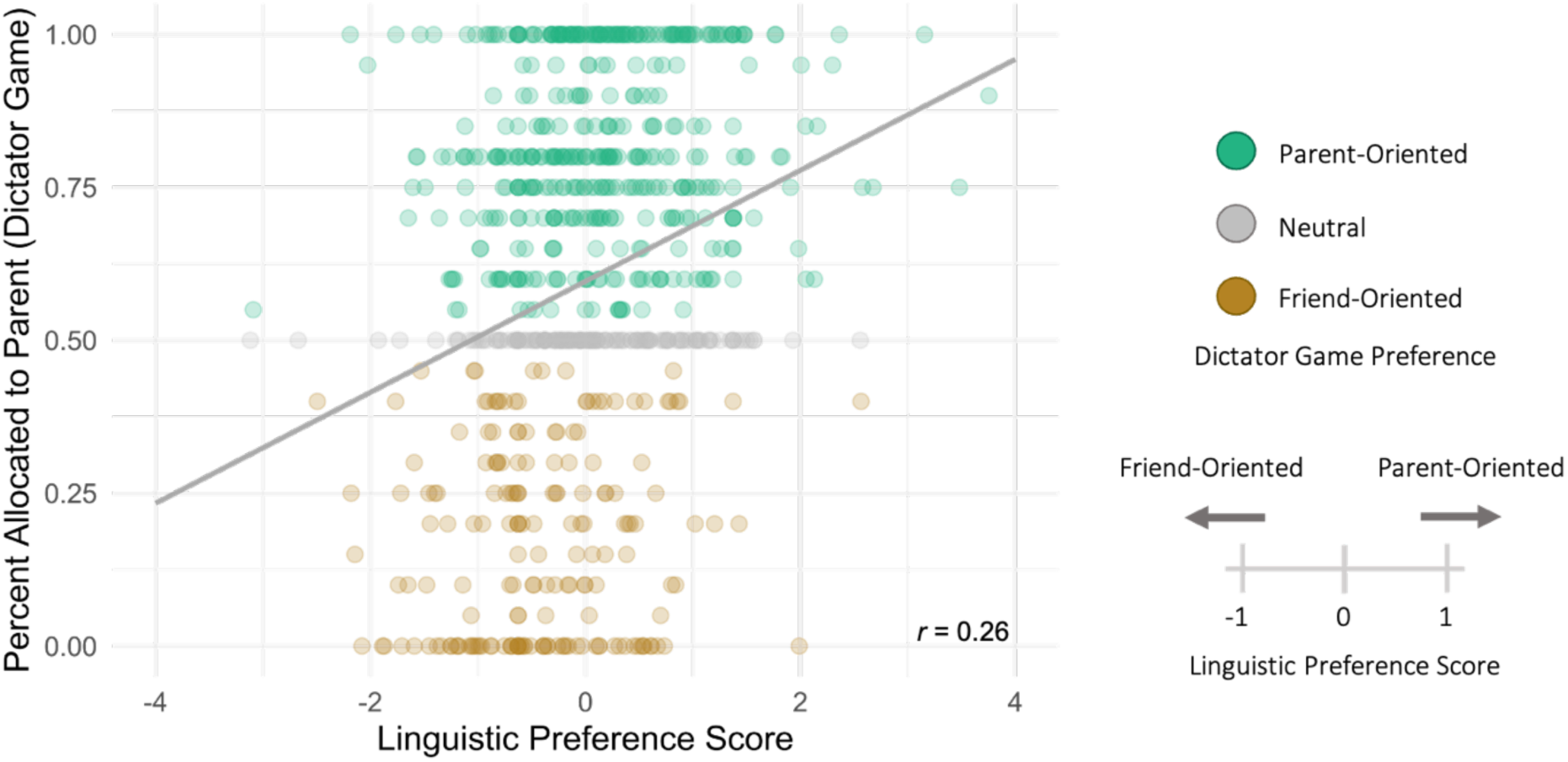
Linguistic preference scores directly predict social decision preferences on a novel decision task (dictator game). *Note.* Linguistic preference scores refer to model implied preferences between a parent and friend based on written text data; relatively greater values indicate a parent-over-friend preference, relatively lower values indicate a friend-over-parent preference. The Y-axis depicts the percentage of a hypothetical $100 endowment allocated to one’s parent (rather than one’s friend); greater values thus indicate stronger parent-over-friend preferences in this task, whereas lower values indicate the opposite. Linguistic preference scores were calculated with weights trained on the Word2Vec-based linguistic representations. Figure 3-1 of the Extended Data shows the relationship of the linguistic preference scores with social loss aversion and relationship quality.

#### Social Decision Preferences are Encoded in Neural Representations of Close Others

After examining differences in parent and friend representations, we next tested whether social decision preferences involving parents and friends were embedded in their neural representations. As noted in the Methods, we estimated trial-level choice preferences between parents and friends using hierarchical Bayesian modeling. The model term that captures choice preferences was allowed to vary randomly across participants, allowing us to model the ensuing variability in preferences with participant-level data about neural representations. Thus, we entered representational similarities between self and other (parent or friend) for each of the six ROIs into four separate models (bilateral ROIs were included in a single model).

For monetary outcomes, we found that greater self-parent similarity in neural representations in the dorsomedial prefrontal cortex (dmPFC) was robustly associated with a greater tendency to favor parents over friends (Table 2). Greater self-friend similarity in the right NAcc was robustly associated with a greater likelihood of favoring friends (Table 3). Curiously, greater self-friend *dis*similarity in the left NAcc was modestly associated with a greater likelihood of favoring friends. For social outcomes, we observed modest evidence that greater self-friend *dis*similarity in mPFC was associated with a greater propensity to favor friends over parents on the task (Table 2). Greater self-parent similarity in the right NAcc was also modestly associated with an increased propensity to favor parents over friends (Table 3). To further ensure that results were not driven by decision-level task features, the models in Tables 2-3 were re-run with all task features replacing reward ratio (discounted reward value, non-discounted reward value, discounted reward time, non-discounted reward time). The results did not appreciably differ when controlling for these other decision-level features.

**Table 2.** Modeling social decision preferences as a function of the overlap of neural representations of self and parent, and self and friend, in dmPFC and mPFC.

|  | Monetary Outcomes |  | Social Outcomes |  |
| --- | --- | --- | --- | --- |
|  | dmPFC | mPFC | dmPFC | mPFC |
| Intercept | **-2.49 [-3.05, -1.93]** | **-2.52 [-3.07, -1.95]** | **-0.92 [-1.33, -0.50]** | **-0.92 [-1.32, -0.50]** |
| Condition | -0.08 [-0.57, 0.44] | -0.06 [-0.63, 0.48] | -0.27 [-0.81, 0.21] | -0.28 [-0.78, 0.19] |
| Reward ratio | -2.96 [-3.56, -2.39] | -2.94 [-3.52, -2.34] | -1.29 [-1.60, -0.93] | -1.28 [-1.62, -0.96] |
| Self-parent neural dissimilarity | 0.45 [0.01, 0.90] | 0.31 [-0.14, 0.80] | 0.02 [-0.34, 0.38] | 0.09 [-0.28, 0.45] |
| Self-friend neural dissimilarity | -0.22 [-0.65, 0.23] | 0.17 [-0.31, 0.62] | 0.01 [-0.36, 0.37] | 0.33 [-0.04, 0.66] |
| Condition x Self-parent neural dissimilarity | **-0.98 [-1.51, -0.44]** | -0.26 [-0.81, 0.34] | -0.23 [-0.78, 0.28] | -0.17 [-0.70, 0.36] |
| Condition x Self-friend neural dissimilarity | -0.07 [-0.58, 0.45] | -0.24 [-0.82, 0.35] | -0.28 [-0.79, 0.26] | **-0.58 [-1.11, -0.07]** |
| LOOIC (SE) | 1322.4 (46.0) | 1325.0 (46.1) | 1703.5 (43.6) | 1704.2 (43.5) |
| Percent Pareto K < 0.7 | 95.5% | 95.5% | 99.8% | 99.7% |

Note. Coefficients are on a logit scale and reflect means from a posterior distribution. ‘Condition’ indicates when parent or friend outcomes were associated with the discounting or delay options and encodes social decision preferences (positive values indicate a parent-over-friend preference, negative values indicate a friend-over-parent preference). Reward ratio reflects the division of the non-discounting option over the discounting option. Self-parent and self-friend neural dissimilarities refer to the distance between multivoxel neural patterns for oneself and a parent or friend, extracted from the brain region listed in the column heading. Neural dissimilarities were calculated via Euclidean distance. Values in brackets reflect 89% highest density credible intervals drawn around the posterior distribution. Interactions reflect associations between neural dissimilarities and social decision preferences. Coefficients where the HDI partially overlaps with ROPE (modest or moderate evidence depending on whether the interval contained 0) are italicized; coefficients where the HDI does not overlap with ROPE (strong evidence) are bolded and italicized. Leave One Out Information Criterion (LOOIC) is presented as a value of fit for each model, along with its standard errors in parentheses. Lower
LOOIC values reflect better fit. The percentage of Pareto K distribution values exceeding a 0.7 cut-off is provided as a diagnostic for gauging the reliability of LOOIC values.

**Table 3.**
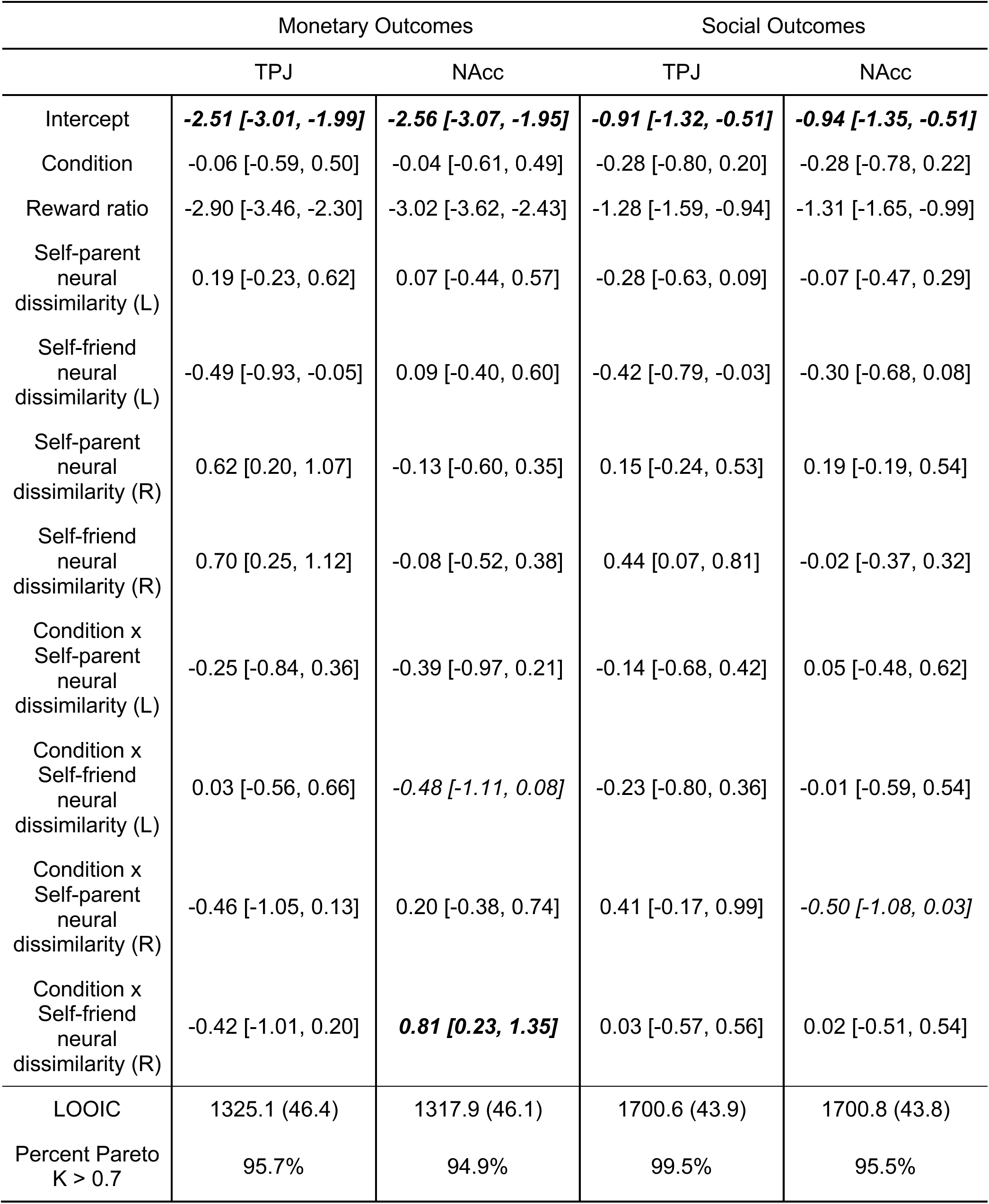
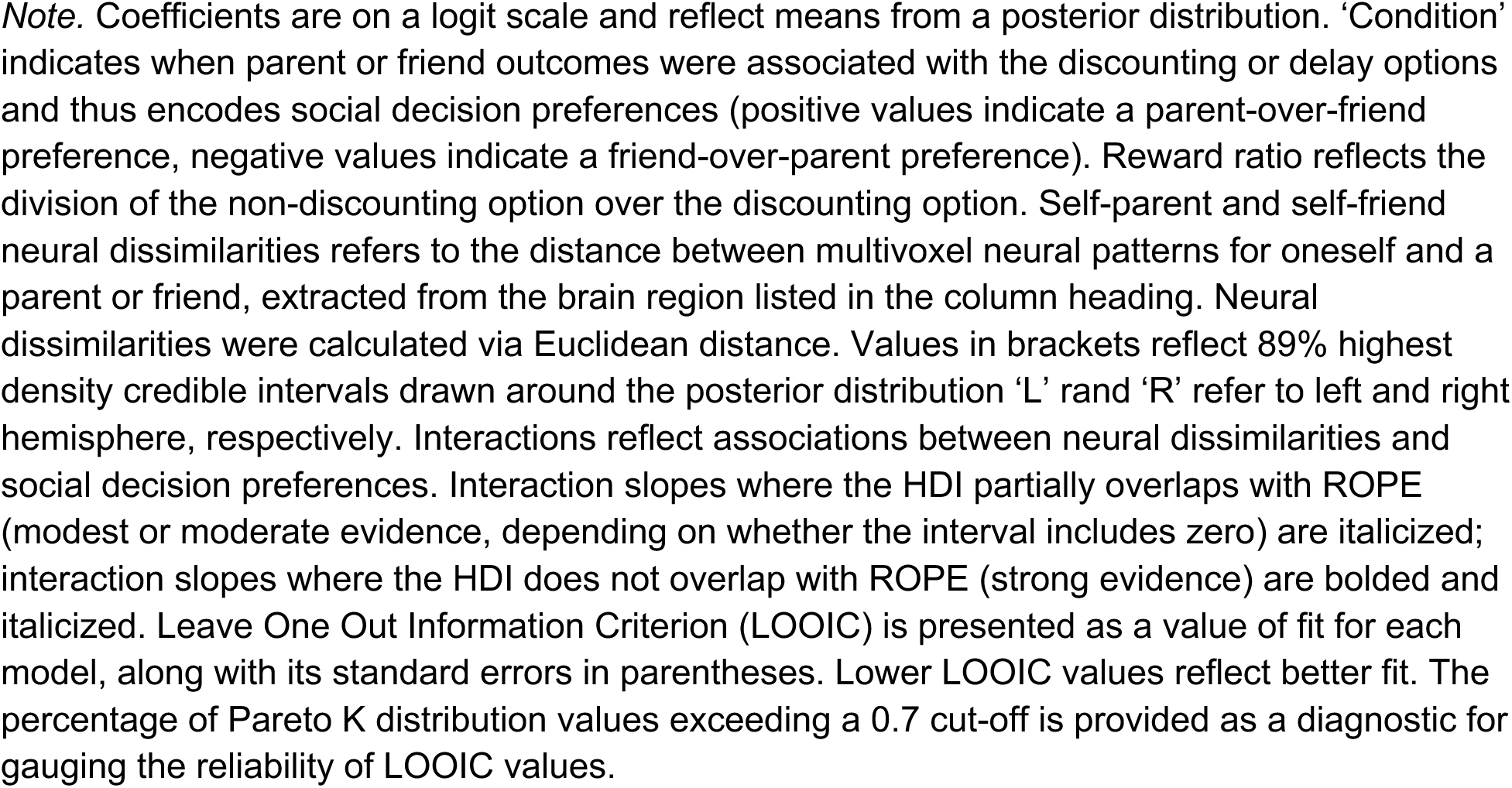
Modeling social decision preferences as a function of the overlap of neural representations of self and parent, and self and friend, bilaterally in the TPJ and NAcc.

|  | Monetary Outcomes |  | Social Outcomes |  |
| --- | --- | --- | --- | --- |
|  | TPJ | NAcc | TPJ | NAcc |
| Intercept | **-2.51 [-3.01, -1.99]** | **-2.56 [-3.07, -1.95]** | **-0.91 [-1.32, -0.51]** | **-0.94 [-1.35, -0.51]** |
| Condition | -0.06 [-0.59, 0.50] | -0.04 [-0.61, 0.49] | -0.28 [-0.80, 0.20] | -0.28 [-0.78, 0.22] |
| Reward ratio | -2.90 [-3.46, -2.30] | -3.02 [-3.62, -2.43] | -1.28 [-1.59, -0.94] | -1.31 [-1.65, -0.99] |
| Self-parent neural dissimilarity (L) | 0.19 [-0.23, 0.62] | 0.07 [-0.44, 0.57] | -0.28 [-0.63, 0.09] | -0.07 [-0.47, 0.29] |
| Self-friend neural dissimilarity (L) | -0.49 [-0.93, -0.05] | 0.09 [-0.40, 0.60] | -0.42 [-0.79, -0.03] | -0.30 [-0.68, 0.08] |
| Self-parent neural dissimilarity (R) | 0.62 [0.20, 1.07] | -0.13 [-0.60, 0.35] | 0.15 [-0.24, 0.53] | 0.19 [-0.19, 0.54] |
| Self-friend neural dissimilarity (R) | 0.70 [0.25, 1.12] | -0.08 [-0.52, 0.38] | 0.44 [0.07, 0.81] | -0.02 [-0.37, 0.32] |
| Condition x Self-parent neural dissimilarity (L) | -0.25 [-0.84, 0.36] | -0.39 [-0.97, 0.21] | -0.14 [-0.68, 0.42] | 0.05 [-0.48, 0.62] |
| Condition x Self-friend neural dissimilarity (L) | 0.03 [-0.56, 0.66] | *-0.48 [-1.11, 0.08]* | -0.23 [-0.80, 0.36] | -0.01 [-0.59, 0.54] |
| Condition x Self-parent neural dissimilarity (R) | -0.46 [-1.05, 0.13] | 0.20 [-0.38, 0.74] | 0.41 [-0.17, 0.99] | *-0.50 [-1.08, 0.03]* |
| Condition x Self-friend neural dissimilarity (R) | -0.42 [-1.01, 0.20] | **0.81 [0.23, 1.35]** | 0.03 [-0.57, 0.56] | 0.02 [-0.51, 0.54] |
| LOOIC | 1325.1 (46.4) | 1317.9 (46.1) | 1700.6 (43.9) | 1700.8 (43.8) |
| Percent Pareto K > 0.7 | 95.7% | 94.9% | 99.5% | 95.5% |

While these results descriptively indicate that social decision preferences are embedded in mental representations of known others, they cannot speak to the out-of-sample predictive power of neural activity measured when thinking about oneself and close others. To address this concern, we conducted a basic predictive modeling analysis, *post hoc,* to verify whether a predictive model using neural RDM cells could predict significantly more variance than a baseline model that only included task-related features. Using leave-one-subject-out cross validation, we employed the glmnet package in R to train a regularized logistic regression model to predict trial-level choices as a function of the condition and reward ratio terms, the neural RDM cells (one model per brain region, averaged across hemispheres for parsimony where relevant), and the interaction between the neural RDM cells and condition.

Imbalanced cases (discount, no discount choices) were handled by assigning weights to each case proportional to the imbalance. Model-predicted choices and observed choices were used to compute a class-weighted composite F1 score for each model (baseline model, baseline model with neural data added). This F1 score assesses joint performance on both the positive (discount) and negative (no discount) cases while taking class imbalance into account. Differences in F1 scores were taken between the baseline model and each of the neural models (Δ_F1_). To perform a significance test of these Δ_F1_ values, we bootstrapped the model output (observed cases, predicted cases) and re-computed the F1 scores for all models, as well as the difference between each neural model and the baseline model, for each of 5,000 iterations, thereby creating an empirical sampling distribution of the Δ_F1_ scores. The results of these analyses are presented in Table 4.

**Table 4.** Predictive models using data from neural RDM cells outperform baseline models that only rely on task data.

| Model | Monetary Outcome | Social Outcome |
| --- | --- | --- |
| Baseline (condition, reward ratio) | 0.589 | 0.553 |
| dmPFC (condition, reward ratio, self-other dmPFC RDM cells, RDM cells x condition interactions) | 0.641<br>$\Delta_{F1} = .052 [0.032, 0.071]$ | 0.562<br>$\Delta_{F1} = .009 [-0.003, 0.023]$ |
| mPFC (condition, reward ratio, self-other mPFC RDM cells, RDM cells x condition interactions) | 0.614<br>$\Delta_{F1} = .025 [0.007, 0.035]$ | 0.584<br>$\Delta_{F1} = .031 [0.013, 0.048]$ |
| NAcc (condition, reward ratio, self-other NAcc RDM cells, RDM cells x condition interactions) | 0.607<br>$\Delta_{F1} = .018 [-0.001, 0.032]$ | 0.578<br>$\Delta_{F1} = .025 [0.001, 0.032]$ |
| TPJ (condition, reward ratio, self-other TPJ RDM cells, RDM cells x condition interactions) | 0.647<br>$\Delta_{F1} = .058 [0.036, 0.079]$ | 0.600<br>$\Delta_{F1} = .047 [0.023, 0.068]$ |

*Note.* The first row in the table lists F1 scores for baseline models predicting choices regarding monetary and social outcomes with task features only (condition, reward ratio). From the second row onward, each row lists the F1 score and a $\Delta_{F1}$ value for a model that was trained with data from the neural RDM cells for a given ROI. Values in brackets refer to 95% bootstrapped confidence intervals for the difference between the F1 score for the model in question and the baseline model. ‘dmPFC’ refers to dorsomedial prefrontal cortex. ‘mPFC’ refers to medial prefrontal cortex. ‘TPJ’ refers to temporoparietal junction. ‘NAcc’ refers to nucleus accumbens.

As shown in Table 4, every model that included interactions between the condition variable and data from neural RDMs evinced a better composite F1 than the baseline (non-neural) model. The added predictive value of models containing neural data was modest but consistent, and amounted to increases in F1 scores of approximately 10% in some cases. In the majority (75%) of cases, the bootstrapped CIs did not overlap with 0, indicating a statistically significant improvement in model performance when incorporating the brain data. In 25% of cases, we did not find evidence to reject the null hypothesis that the F1 difference with vs. without including the neural data is 0; this could be due to several factors, including differences between the Bayesian approach used in our main analyses vs. this bootstrapping approach, differences stemming from collapsing across hemispheres in the bootstrapping analyses but not the main (Bayesian) analyses, and/or the fact that this predictive modeling approach is penalizing model predictions to ensure generalizability, which could come at the expense of understanding small but meaningful associations between variables. That said, in the majority of cases, incorporating the neural data significantly improved model predictions of participants’ choices, over and above what was achieved when basing predictions only on task features.

### Social decision preferences are encoded in linguistic representations of close others

#### Social Decision Preferences are Encoded in Linguistic Representations of Close Others

The average leave-one-subsample-out classification accuracies for each of the three candidate models when engineering the linguistic were roughly comparable (SVC: Mean = 0.554, SD =.04, Range = [0.489, 0.585], RRC: Mean = 0.566, SD =.06, Range = [0.471, 0.622], RLR: Mean = 0.562, SD =.06, Range = [0.480, 0.622]). We decided to select weights from the SVC model because it had relatively high accuracy and the lowest performance variance. Notably, when taking into account the baseline rate of parent-over-friend preferences in each of these samples, the classification accuracies for each sample were statistically better than chance when considering sample size. That social decision behavior could be successfully decoded from written text was the first piece of evidence that linguistic representations of close others contain motivationally relevant information about participants’ preferences.

To follow this up, we tested whether social decision preferences in novel data with different assessments of social choice preferences could be predicted as function of linguistic preference scores – or, one’s social decision preference implied by the model. Model weights from the linguistic signature were thus applied to novel data to estimate associations with social decision preferences in a one-shot dictator task in which participants allocated a finite sum of hypothetical money between their parent and friend (Figure 3). Linguistic preference scores were robustly associated with preferences on this task (*r* = 0.26, 89% HDI = [0.21, 0.31]). We then observed modest evidence that linguistic preference scores exhibited a small association with social decision preferences between parents and friends in the multi-trial risky decision task (*b* = 0.02 [-0.06, 0.11]). Preference scores derived from our linguistic signature were robustly associated with choice preferences between parents and friends in a multi-trial delay discounting task, when both monetary (*b* = 0.94, [0.72, 1.20]) and social (*b* = 1.14, [0.91, 1.39]) outcomes were at stake. Finally, we were able to show that linguistic preference generalized such that they were robustly associated with *attitudes* about one’s relationship with their parents (*r_parent_* = 0.23, [0.19, 0.27]). Evidence for null effects was observed for the relationship between preference scores and friend relationship quality, however (*r_friend_* = 0.00, [-0.04, 0.04]). These findings are visualized in Figure 3-1.

### Linguistic and Neural Representations are Correlated, and Jointly Predict Social Decision Preferences

The preceding findings raise the question of whether having both a strong linguistic preference score and extensive overlap between neural representations of oneself and a given other is associated with a stronger social decision preference for that individual. In other words, if someone’s representations of others were ‘biased’ towards a particular other (e.g., one’s friend rather than one’s parents) in *both* neural and linguistic data, would we also observe even stronger social decision preferences? Such a pattern of results would have several implications for what kinds of information are stored in neural and linguistic representations, and how they are linked with each other and behavior.

The degree of correspondence between linguistic preference scores and neural similarity is shown in Figure 4 and Table 5. We found that preference scores implied by the linguistic signature were correlated with similarity between self and parent neural representations in the anticipated direction for three of the six brain regions (mPFC, R NAcc, L TPJ). Neural similarity between self and friend was correlated with linguistic preferences in the anticipated direction for five of the six brain regions (dmPFC, mPFC, L NAcc, L TPJ and R TPJ).

**Figure 4.**
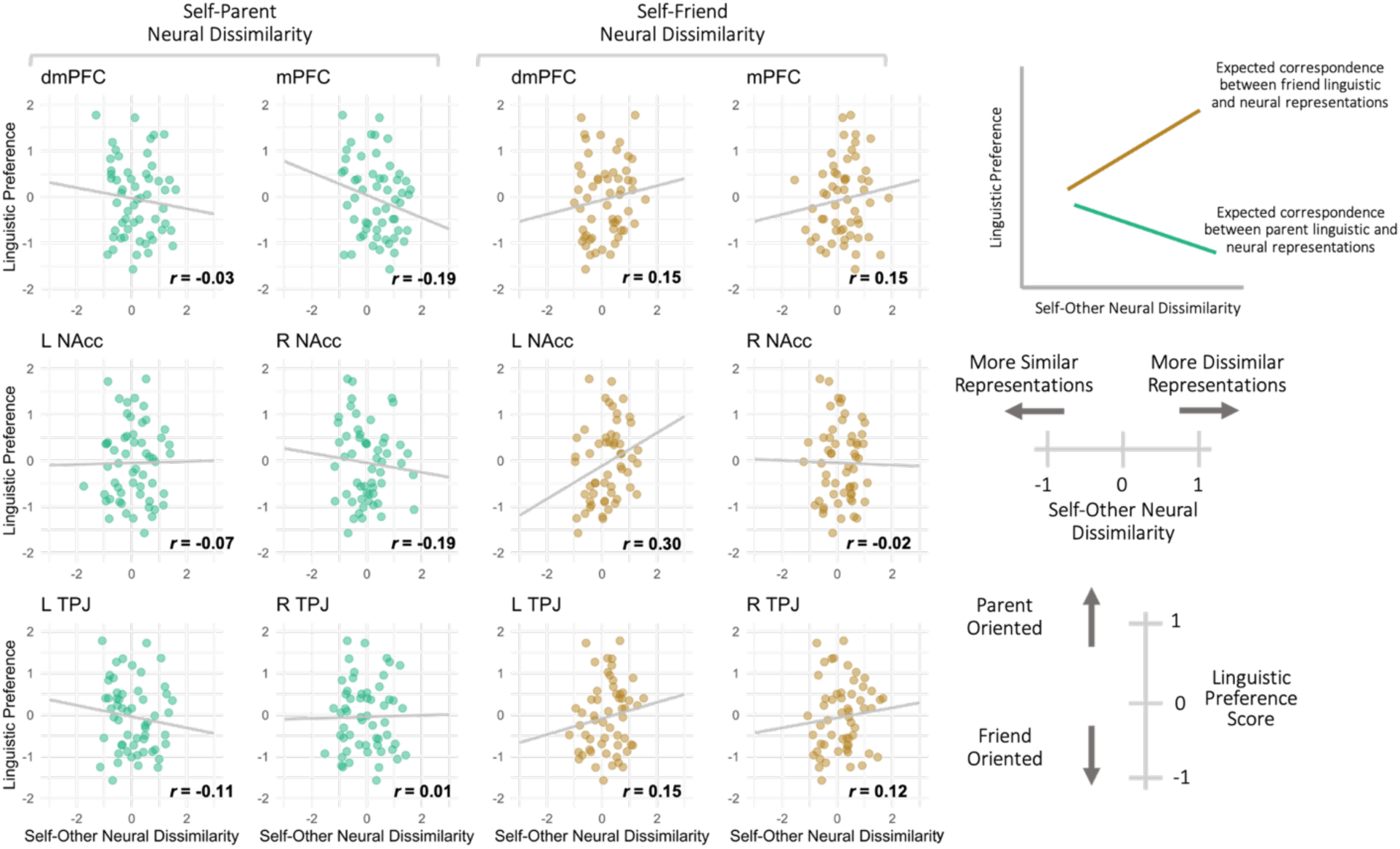
Neural and linguistic representations of parents and friends are correlated in the TPJ, NAcc, and dmPFC. *Note.* Linguistic preference scores refer to model implied social decision preferences between a parent and friend based on written text data; relatively greater values indicate a parent-over-friend preference, relatively lower values indicate a friend-over-parent preference. The Y-axis depicts neural dissimilarity between self and other (parent, friend) representations across six neural ROIs. Linguistic preference scores were calculated with weights trained on Word2Vec-based linguistic representations.

**Table 5.**
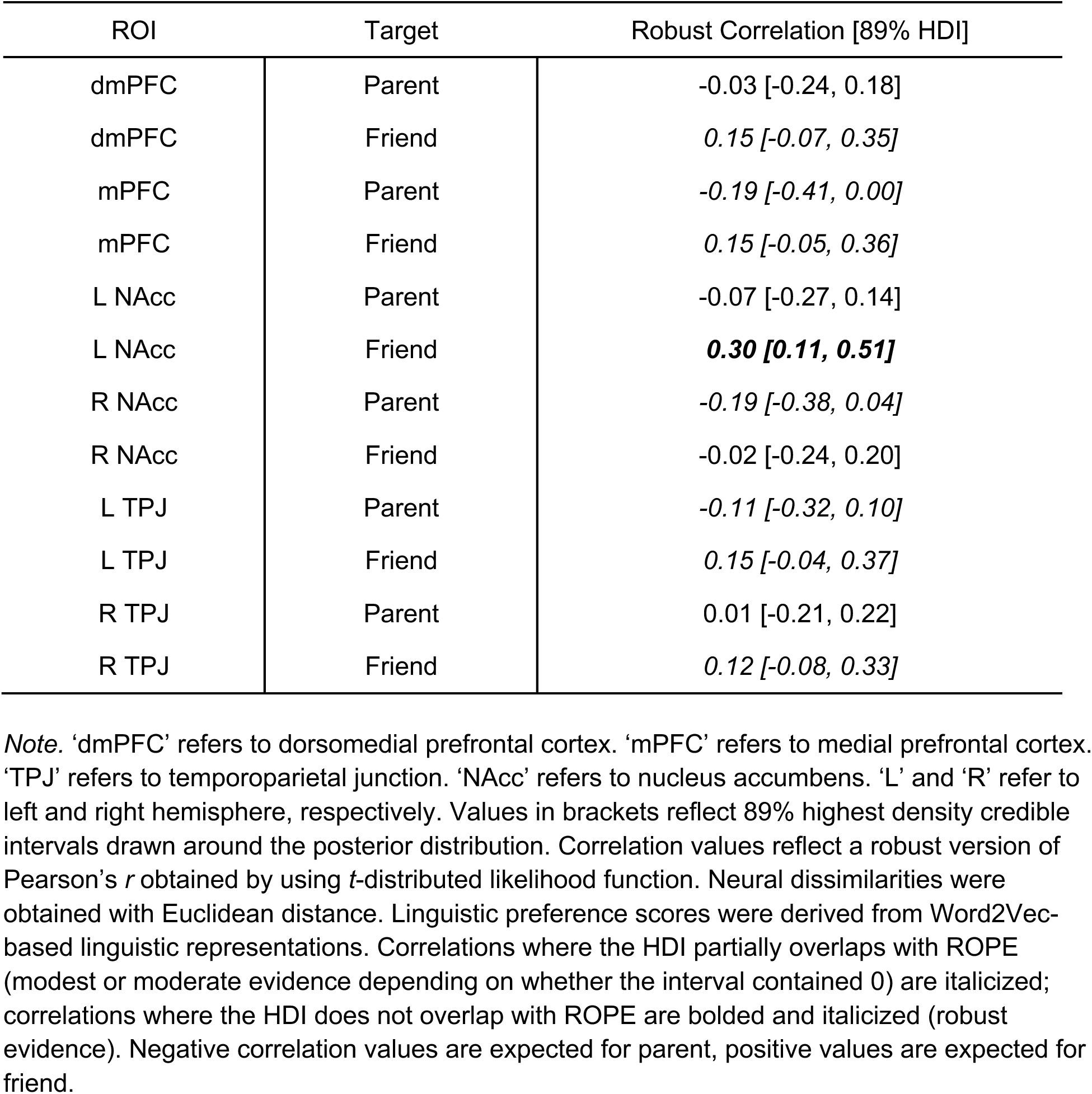
Correlations between linguistic preference scores and neural representational dissimilarities between self and other across two close others (targets) and six ROIs.

We then examined whether linguistic preference scores moderated the link between neural representational similarity and choice preferences on the discounting task (Figure 5, Tables 6-1, 6-2). The logic behind this analysis is that linguistic representations and neural representations obtained with the methods used here may capture partially distinct aspects of one’s mental representations of others. Thus, combining these different modalities could afford greater sensitivity for detecting associations between one’s mental representations of others and behavior. In the monetary outcome domain, a relationship in the non-anticipated direction for the L NAcc was observed with linguistic preference scores and neural dissimilarity between self and friend representations. No other effects met our evidentiary criteria or appeared close to meeting them in the monetary outcome domain. In the social outcome domain, we found that linguistic preference scores interacted with self and parent neural dissimilarity in the mPFC and R NAcc to promote stronger social decision preferences for parents if neural similarity was high and linguistic preference scores were high. A similar finding with self and friend neural dissimilarity in the left TPJ was indicated by examining the posterior distribution but did not quite meet our evidentiary criteria.

**Figure 5.**
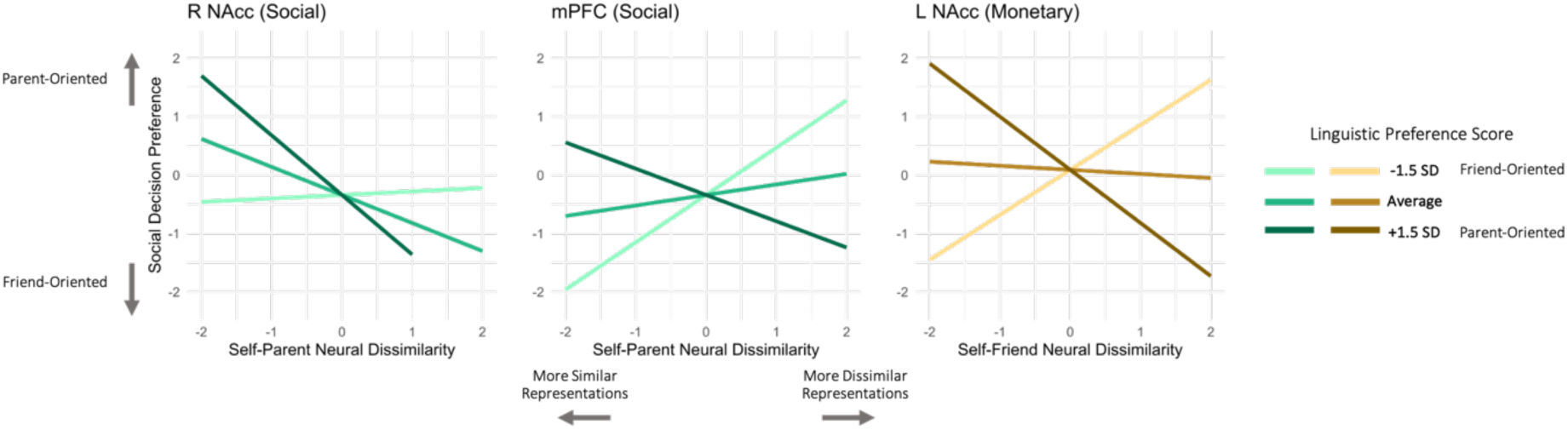
Correspondence between linguistic and neural representations in three ROIs is associated with stronger decision preferences. *Note.* Linguistic preference scores refer to model implied social decision preferences between a parent and friend based on written text data; relatively greater values indicate a parent-over-friend preference, relatively lower values indicate a friend-over-parent preference. The Y-axis represents choice preferences between parent and friend. The X-axis depicts neural representational dissimilarity between the self and a target other (parent, friend) for a given ROI. The relevant ROI is listed at the top of the top of plot, along with the outcome domain of the decision task (monetary or social). Linguistic preference scores were calculated with weights trained on the Word2Vec-based linguistic representations.

**Table 6.**
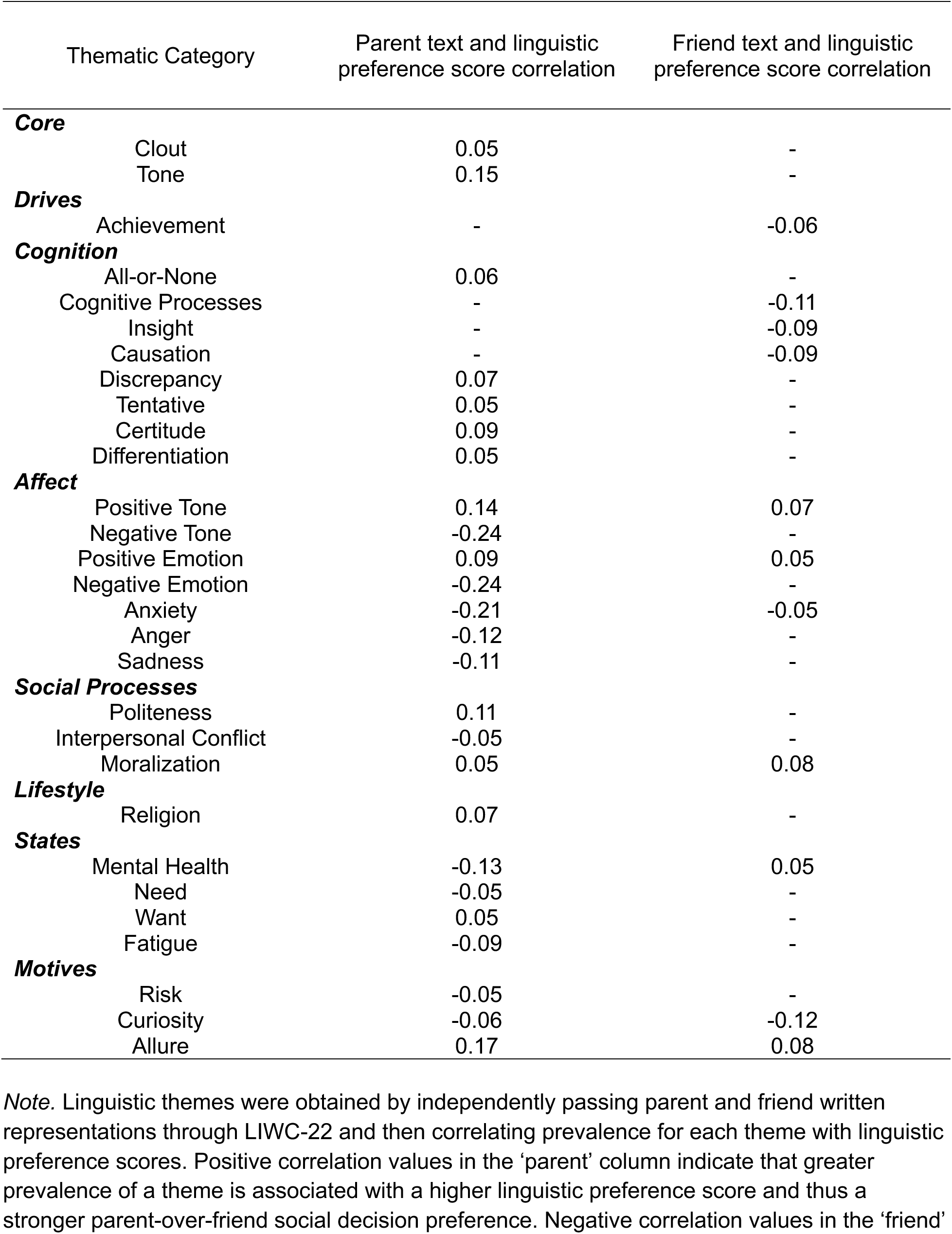
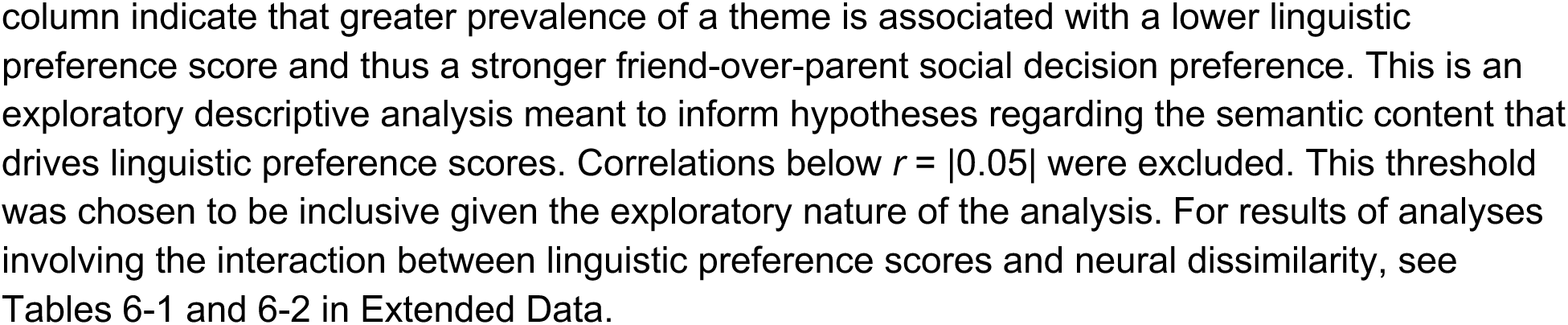
Linguistic themes from parent and friend linguistic representations most strongly correlated with linguistic preference scores.

| Thematic Category | Parent text and linguistic preference score correlation | Friend text and linguistic preference score correlation |
| --- | --- | --- |
| **Core** |  |  |
| Clout | 0.05 | - |
| Tone | 0.15 | - |
| **Drives** |  |  |
| Achievement | - | -0.06 |
| **Cognition** |  |  |
| All-or-None | 0.06 | - |
| Cognitive Processes | - | -0.11 |
| Insight | - | -0.09 |
| Causation | - | -0.09 |
| Discrepancy | 0.07 | - |
| Tentative | 0.05 | - |
| Certitude | 0.09 | - |
| Differentiation | 0.05 | - |
| **Affect** |  |  |
| Positive Tone | 0.14 | 0.07 |
| Negative Tone | -0.24 | - |
| Positive Emotion | 0.09 | 0.05 |
| Negative Emotion | -0.24 | - |
| Anxiety | -0.21 | -0.05 |
| Anger | -0.12 | - |
| Sadness | -0.11 | - |
| **Social Processes** |  |  |
| Politeness | 0.11 | - |
| Interpersonal Conflict | -0.05 | - |
| Moralization | 0.05 | 0.08 |
| **Lifestyle** |  |  |
| Religion | 0.07 | - |
| **States** |  |  |
| Mental Health | -0.13 | 0.05 |
| Need | -0.05 | - |
| Want | 0.05 | - |
| Fatigue | -0.09 | - |
| **Motives** |  |  |
| Risk | -0.05 | - |
| Curiosity | -0.06 | -0.12 |
| Allure | 0.17 | 0.08 |

*Note.* Linguistic themes were obtained by independently passing parent and friend written representations through LIWC-22 and then correlating prevalence for each theme with linguistic preference scores. Positive correlation values in the 'parent' column indicate that greater prevalence of a theme is associated with a higher linguistic preference score and thus a stronger parent-over-friend social decision preference. Negative correlation values in the 'friend'
column indicate that greater prevalence of a theme is associated with a lower linguistic preference score and thus a stronger friend-over-parent social decision preference. This is an exploratory descriptive analysis meant to inform hypotheses regarding the semantic content that drives linguistic preference scores. Correlations below $r = |0.05|$ were excluded. This threshold was chosen to be inclusive given the exploratory nature of the analysis. For results of analyses involving the interaction between linguistic preference scores and neural dissimilarity, see Tables 6-1 and 6-2 in Extended Data.

While these results indicate some degree of informational correspondence between neural and linguistic representations and subsequently suggest neural representations driving behavior are comprised of semantic information, it remained an open question about what kinds of semantic content are indexed in these representations.

To unpack the statistical relationship between neural representations and linguistic preference scores we conducted an exploratory, descriptive analysis using the most recent version of the Linguistic Inquiry and Word Count (LIWC-22) software (Boyd et al., 2022) to identify the semantic themes that drive linguistic preference scores and then correlated them with neural representations. We did this in two steps.

First, we passed the written text data of Study 2 participants through LIWC-22 and extracted the prevalence of the following thematic categories (individual facets in parentheses): core summary themes (clout, analytical thinking, authentic, emotional tone), drives (affiliation, achievement, power), cognition (all-or-none, cognitive processes, insight, causation, discrepancy, tentative, certitude, differentiation), affect (positive tone, negative tone, positive emotion, negative emotion, anxiety, anger, sadness), social processes (prosocial behavior, politeness, interpersonal conflict, moralization, communication), culture (politics, ethnicity, technology), lifestyle (leisure, home, work, money, religion), states (mental health, need, want, acquire, lack, fulfilled, fatigue), and motives (reward, risk, curiosity, allure). The prevalence of each of these themes in parent and friend text was then separately correlated with linguistic preference scores. We kept themes that showed a correlation of *r* > |.05|. This threshold was chosen because it roughly corresponded to the threshold for statistical significance in this subsample and because it minimized the risk of falsely excluding an interesting or meaningful theme. Results indicate that affective themes (e.g., risk, want, fatigue) in parent text are the primary drivers of linguistic preference scores; cognitive and state-based themes (e.g., causation, mental health) are ostensibly the primary drivers of linguistic preference scores for friend text (Table 6). Notably, themes in parent text tended to covary more with linguistic preference scores than those in friend text insofar that more parent text themes evinced correlations that met our threshold.

Second, we took the themes identified in Table 6 and extracted them from Study 1 participants’ text data, and then correlated their prevalence in that data with self-other neural similarity from ROIs that showed the strongest relationships with linguistic preference scores (Parent: R NAcc, mPFC, dmPFC, L TPJ; Friend: L NAcc, mPFC, dmPFC, L TPJ; see Figures 4-5). These correlations are depicted as Cleveland plots in Figures 6-7. The most consistently strong themes for parents and friends were identified by taking the magnitude (absolute value) of each theme’s correlation with neural data and averaging it across the ROIs. These are listed in the top right corner of Figures 6-7. Because the purpose of this analysis was meant to descriptively unpack what is happening in our sample and help generate future research, we did not conduct inferential statistics on the estimates. Affective and state-based themes were most consistently associated with self-parent representational overlap across ROIs, whereas cognitive themes were most consistently associated with self-friend representational overlap across ROIs. Because of potential negations in the text data (e.g., ‘I do not feel happy’) we encourage readers to primarily interpret the magnitude, rather than the sign, of these correlations.

**Figure 6.**
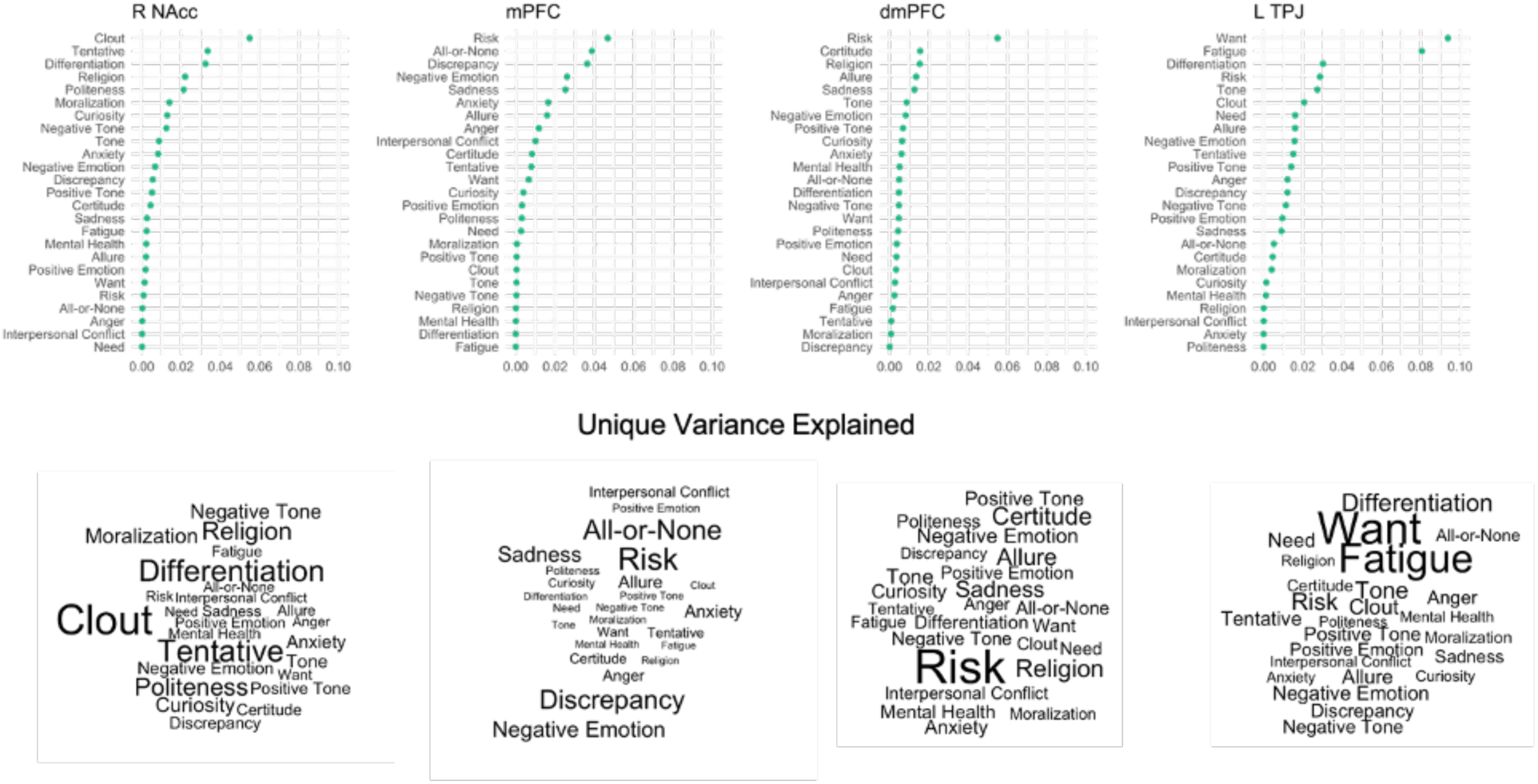
Unpacking the correspondence between linguistic and neural representations of parents: Unique variance explained in self-parent neural dissimilarity by each linguistic theme. *Note.* Linguistic themes analyzed here are those from the ‘parent’ column of Table 6. ROIs analyzed here are those showing the strongest relationship with linguistic preference scores (Figures 4-5). Datapoints represent the squared semi-partial correlation between the prevalence of a linguistic theme in written text data (about parents) and self-parent neural dissimilarity in an ROI (i.e., the unique variance each theme accounts for in the neural data given all other themes). This is an exploratory descriptive analysis meant to understand what semantic content is encoded in self-other neural representations. Pearson’s *r* coefficient was used. Each term in the word cloud corresponds to a linguistic theme, and their sizes are scaled by the unique variance explained.

**Figure 7.**
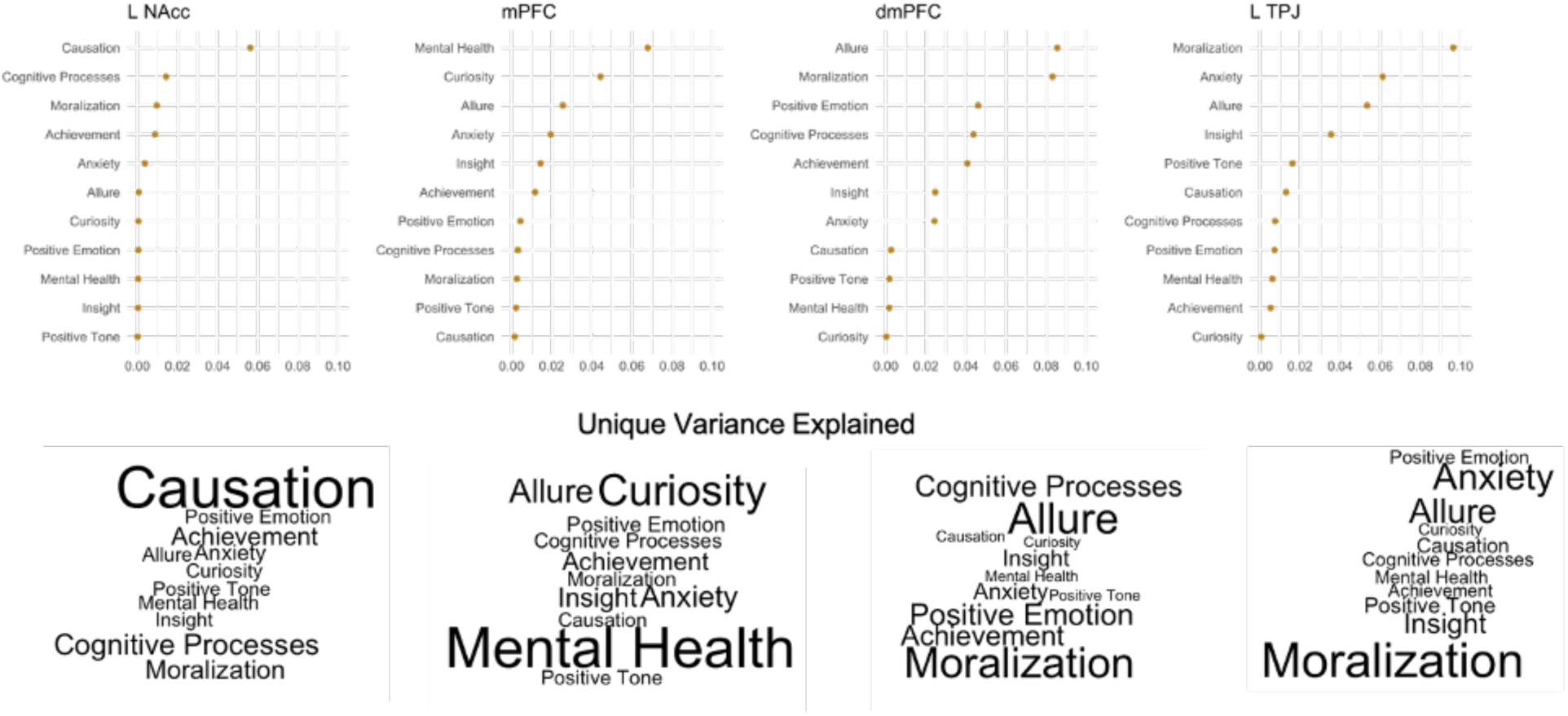
Unpacking the correspondence between linguistic and neural representations of friends: Unique variance explained in self-friend neural dissimilarity by each linguistic theme. *Note.* Linguistic themes analyzed here are those from the ‘friend’ column of Table 6. ROIs analyzed here are those showing the strongest relationship with linguistic preference scores (Figures 4-5). Datapoints represent the squared semi-partial correlation between the prevalence of a linguistic theme in written text data (about friends) and self-friend neural dissimilarity in an ROI (i.e., the unique variance each theme accounts for in the neural data given all other themes). This is an exploratory descriptive analysis meant to understand what semantic content is encoded in self-other neural representations. Pearson’s *r* coefficient was used. Each term in the word cloud corresponds to a linguistic theme, and their sizes are scaled by the unique variance explained.

## Discussion

In the current study, we found evidence that neural and linguistic representations of close others contain information about one’s social decision preferences involving these individuals. We first observed that neural representations of parents and friends are differentially encoded in the brain with respect to the self, such that representations of one’s parents are more similar to representations of oneself in bilateral NAcc and TPJ, whereas representations of one’s friends are more similar to representations of oneself in mPFC. Individual differences in the encoding ‘bias’ of these brain regions were related to and possibly consequential for social decision preferences involving close others, as self-other representational similarities in the NAcc, TPJ, and dmPFC tracked with between-individual social decision preferences.

Linguistically, written memories and descriptors of parents and friends were used to train a linguistic signature of social decision preferences. When applied to new data, the signature was able to predict choice preferences between parents and friends on novel decision tasks and was even related to other relationship characteristics (e.g., relationship quality). Critically, the scores from the linguistic signature and neural data were related, showing correspondence of representations measured with linguistic and neural data. Finally, we found that individual differences in the degree of linguistic-neural correspondence were also predictive of social decision preferences. These findings collectively enrich our understanding of how the mental models that humans create for *specific* others relate to choices regarding those others.

Existing studies on social decision-making predominantly focus on understanding decision behavior in terms of the cognitive heuristics that individuals apply over situation-level features or how individuals weigh the mental states of others (Austerweil et al., 2016; Cole & Bruno Teboul, 2004; Crockett et al., 2017; FeldmanHall & Chang, 2018; Hackel et al., 2017; Kao et al., 2023; Sampaio et al., 2023; Yu et al., 2019). Our findings suggest that an additional driver of social decision preferences stems from how representations of oneself and others are aligned in the brain. We found that self-other similarity in mPFC and dmPFC was associated with choice preferences between a parent and friend. Specifically, greater neural representational overlap between self and parent was associated with a greater likelihood of favoring one’s parent-over-friend, whereas *less* self and friend representational overlap (i.e., greater dissimilarity) was associated with stronger friend-over-parent preferences. The NAcc was also predictive of choice preferences, but the direction of the effect varied based on several factors. These varied patterns, paired with existing psychology and neuroscience literature showing that representations of others are linked to a wide range of cognitive, affective, and semantic processes (Guassi Moreira et al., 2023; Lin & Thornton, 2023; Tamir & Thornton, 2018; Todd & Tamir, 2024), suggest these brain regions could be storing and accessing different types of representational information and/or associated mental processes. Indeed, descriptive patterns in the written text data appear to provide some evidence for this possibility. Variation in the specific brain regions implicated as a function of outcome type (monetary or social) is consistent with the notion that brain regions store and access different kinds of representational information that may be selectively accessed in different decision contexts. That said, we note that similarities in representations of oneself and others in the NAcc were associated with decision preferences across both outcome types. This suggests that representations of familiar others in the NAcc region may shape social decisions regarding those individuals across a wide range of contexts and outcomes, which may be related to the motivational and reward-related process that this region supports. Our pattern of findings overall hint at the idea that ‘encoding biases’ towards one’s parent or friend in the brain regions studied are related to choice preferences implies the possibility of affordances in these representations (de Wit et al., 2017; Kourtis et al., 2018; McMahon & Isik, 2023), such that the way in which information (here, representations of close others) is encoded might facilitate particular behavioral tendencies during decision-making.

Relatedly, our linguistic findings—the prediction of choice preferences from text data and model weights generalizing to other facets of relationships—point towards a similar concept of ‘psychological affordances’. The most readily available mental information when prompted to describe a close other or write down a memory about them (the prompts used to elicit the text data that was used to train the linguistic signature) may also be the most readily available or salient information when making choices that impact said close other. That our decision tasks did not allow for prolonged deliberation between trials meant there was a degree of spontaneity required when responding. Having one’s representations of other people contain behavioral affordances is a potentially efficient facet of the psychological architecture that links mental models with behaviors.

The joint associations of neural and linguistic mental representations with decision preferences here are a particularly interesting facet of investigation with respect to the ability of linguistic information to disambiguate brain-behavior associations. fMRI data cannot always be used to infer the exact operations performed by the brain. However, close correspondence between linguistic and neural data can refine hypotheses about the types of information the brain is using. Here we observed mixed evidence: while linguistic preference scores were broadly correlated with imaging data, high correspondence between linguistic and neural information (e.g., strong correlation between neural similarity and linguistic preference scores) did not always augment social choice preferences. The results of these analyses can be interpreted in different ways. On the one hand, if indices of neural and linguistic representations show high correspondence, it could indicate convergence in the types of information carried by each and make one of the pair redundant when modeling social decision preferences (such as in the case when monetary outcomes were at stake). On the other hand, if a high correlation between linguistic and neural representational information was redundant, then why would it be associated with augmented decision preferences (such as the case with social outcomes)? One explanation could be that having a high linguistic preference score could indicate a shift in the type of content encoded in mental representations in such a way that is possibly more consequential for shaping decision preferences. That this occurs for social but not monetary outcomes *could* mean that decisions involving spending time with others are particularly sensitive to representational content that is most readily accessed in autobiographical content, though this would need to be tested directly. Linguistically, neural representational overlap with both parent and friend relationships was associated with motivationally relevant themes. Parent neural representations were more uniquely linked with state-based themes, whereas friend neural representations evinced stronger relationships with social cognitive and mental health related themes. While the results of these analyses are meant to be descriptive and to inform hypotheses for future research, they broadly suggest that representations of specific familiar others are comprised of distinct psychological content.

While our findings highlight an important association between mental representations and social decision-making, and we hypothesize that mental representations are a driving force in shaping behavior, we must also acknowledge that one’s social choice preferences involving close others may very well sculpt the architecture of mental representations. Contextual factors unrelated to mental representations may also constitute potent forces that shape initial social decision preferences, which then in turn may influence the construction of mental representations. Another possibility is that a bidirectional relationship may exist such that individuals who prioritize their relationships with particular familiar others may develop mental representations of those people that are more similar to their mental representations of themselves, which could subsequently influence their downstream decision-making. Future research should explore this interplay more deeply, potentially using longitudinal designs to examine how relationship dynamics, decision preferences, and mental representations evolve over time. Additionally, qualitative methodologies could provide insights into how individuals navigate these complexities in real-world decision-making scenarios.

Future studies could improve and extend on ours in several ways. First, researchers may consider testing other types of familiar relationship categories, not just close others.

Understanding how mental representations influence behavior is likely best accomplished by examining such associations across the entire spectrum of psychological and social distance between an individual and the people in their social environment. Such efforts may also consider exhaustively characterizing social decision preferences on a wider range of tasks with varied experimental conditions (e.g., including self versus other conditions). Keeping with this spirit, it would also be helpful to conduct similar studies across different populations to assess the generalizability of these findings. Second, longitudinal work is critical for quantifying the temporal dynamics of how representations are formed and updated with experience, as this is critical for making causal claims. Such avenues would benefit from complementary use of intensive longitudinal methods such as ecological momentary assessment, or incorporating developmental populations that contain individuals who are frequently introduced to completely novel social networks (e.g., incoming college students, middle or high schoolers) and thus may serve as ideal populations for pursuing this line of work.

On a related note, given that mental representations of others are not static entities, another benefit of longitudinal work would be to examine how mental representations of others fluctuate over time and how such fluctuations correspond to social decision-making behavior. Additionally, some of the benefits of longitudinal work can also perhaps be attained without multiple sessions, by varying the temporal ordering between social decision and representation-eliciting tasks within a session to measure whether order effects exist and whether the decisions that people make regarding familiar others impact their mental representations of those individuals. Third and last, we believe that future work should incorporate more extensive written text data about close others so as to bolster sensitivity by more extensively sampling one’s mental representations of those individuals. Ideally, future studies utilizing intensive longitudinal methods would incorporate written diaries or essay-like prompts at baseline to gather even more text data about close others, or such studies could ethically obtain written records (e.g., text messages) between participants and close others. Such approaches could also be used to help disentangle the role of mental representations in shaping decision behavior versus how cognitive heuristics such as aversion to inequity, risk, or ambiguity may be differentially deployed when making choices that affect others compared to oneself. Longitudinal approaches may be particularly helpful for this issue because they allow for the use of lagged analyses could help better delineate the relative influence of mental models of others versus differential deployment of cognitive heuristics.

Finally, our study speaks to the importance of studying known close others. This study is another entry in a growing body of work that shows individuals have highly granular behavioral choice tendencies involving other individuals (Brandner et al., 2021; Delton et al., 2023; Karan et al., 2022; Sznycer, 2022). Understanding human behavior requires examining the most relevant and meaningful phenomena in our daily experiences. Knowing *how* individuals represent specific others and in turn use these representational distinctions to guide behavior is critical for understanding much of human behavior as it unfolds outside the laboratory.

## Acknowledgments

Preparation of this research was supported by a National Science Foundation SBE Postdoctoral Research Fellowship to JFGM (award number 2104629) and the UCLA Department of Psychology (CP). We thank Dr. Rick Dale for comments on the linguistic analyses. We thank the members of the Computational Social Neuroscience Laboratory at UCLA for their feedback on study concept and manuscript.

## Extended Data – Figure Captions

**Figure 1-1.**
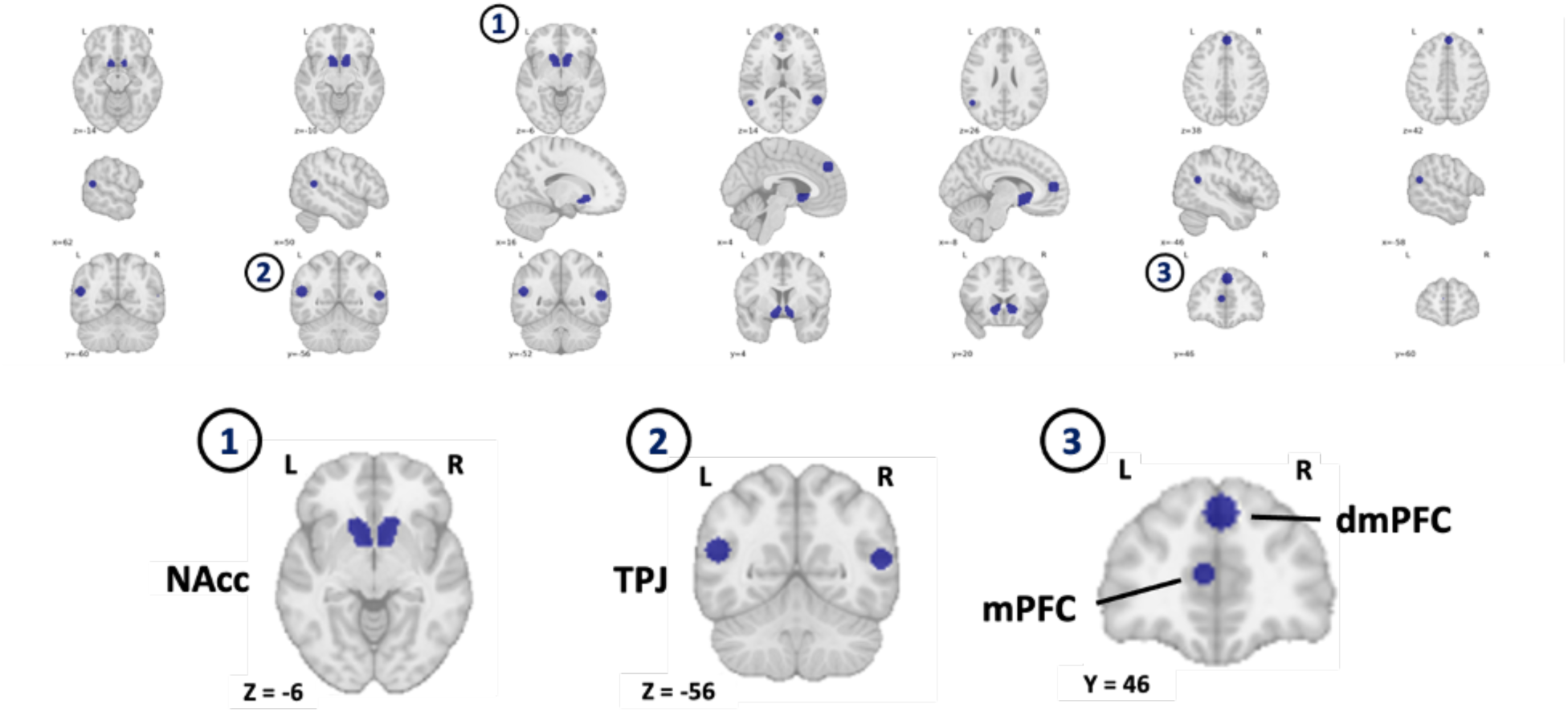
The six masks used for ROI analyses. *Note.* Masks for the mPFC, dmPFC, and bilateral TPJ were created using Neurosynth. Bilateral NAcc masks were created using the Harvard-Oxford probabilistic atlas.

**Figure 2-1.**
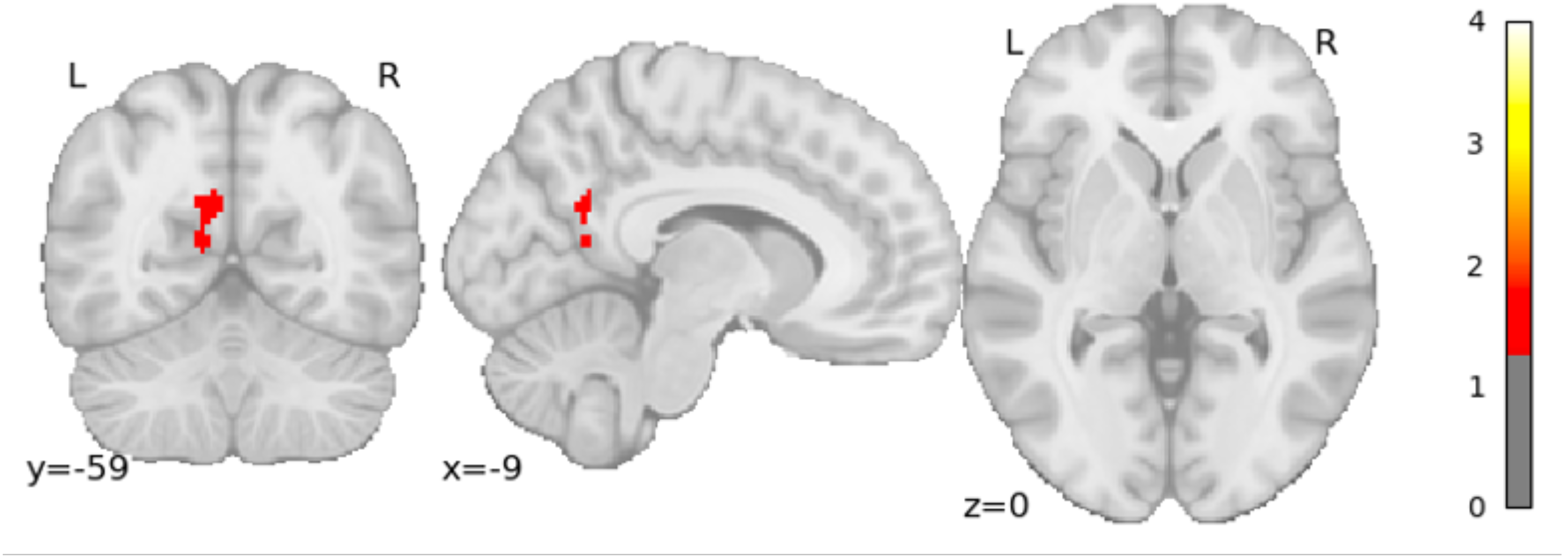
Exploratory searchlight results showing distinct representations of parents and friends. *Note.* Group level analyses were conducted via permutation test (10,000 iterations) using Nilearn’s non_parametric_inference function and TFCE (FWE, *p* <.05). Negative log10 p-values are displayed as cluster intensity values. *P-*values are corrected for multiple comparisons (FWE, *p* <.05). The color bar ranges correspond to the threshold needed to achieve FWE error control (1.3) and the lowest possible negative log10 *p*-value given the number of permutations (4).

**Figure 2-2.**
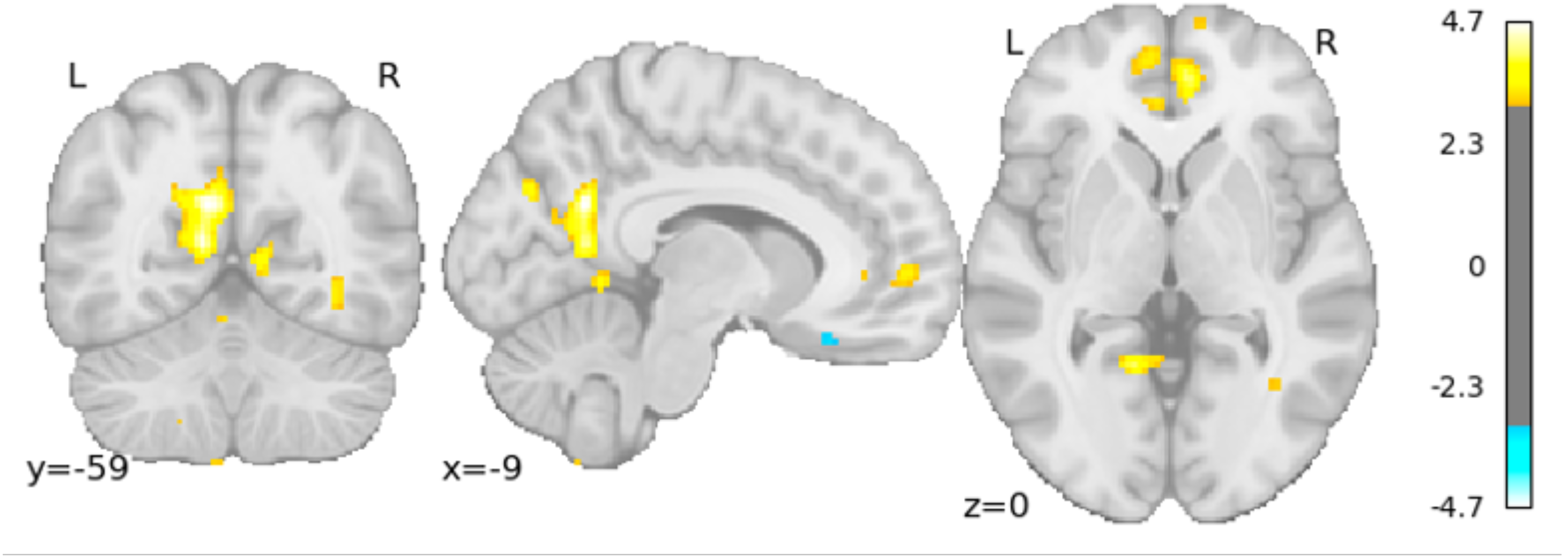
Uncorrected results of the exploratory searchlight analysis probing for distinct representations of parent and friend representations. *Note.* Group level analyses were conducted via permutation test (10,000 iterations) using Nilearn’s non_parametric_inference function. *t*-statistics are displayed for cluster intensity values (thresholded at 3.1). No corrections for multiple comparisons have been made to this image; it is plotted purely for exploratory purposes.

**Figure 2-3.**
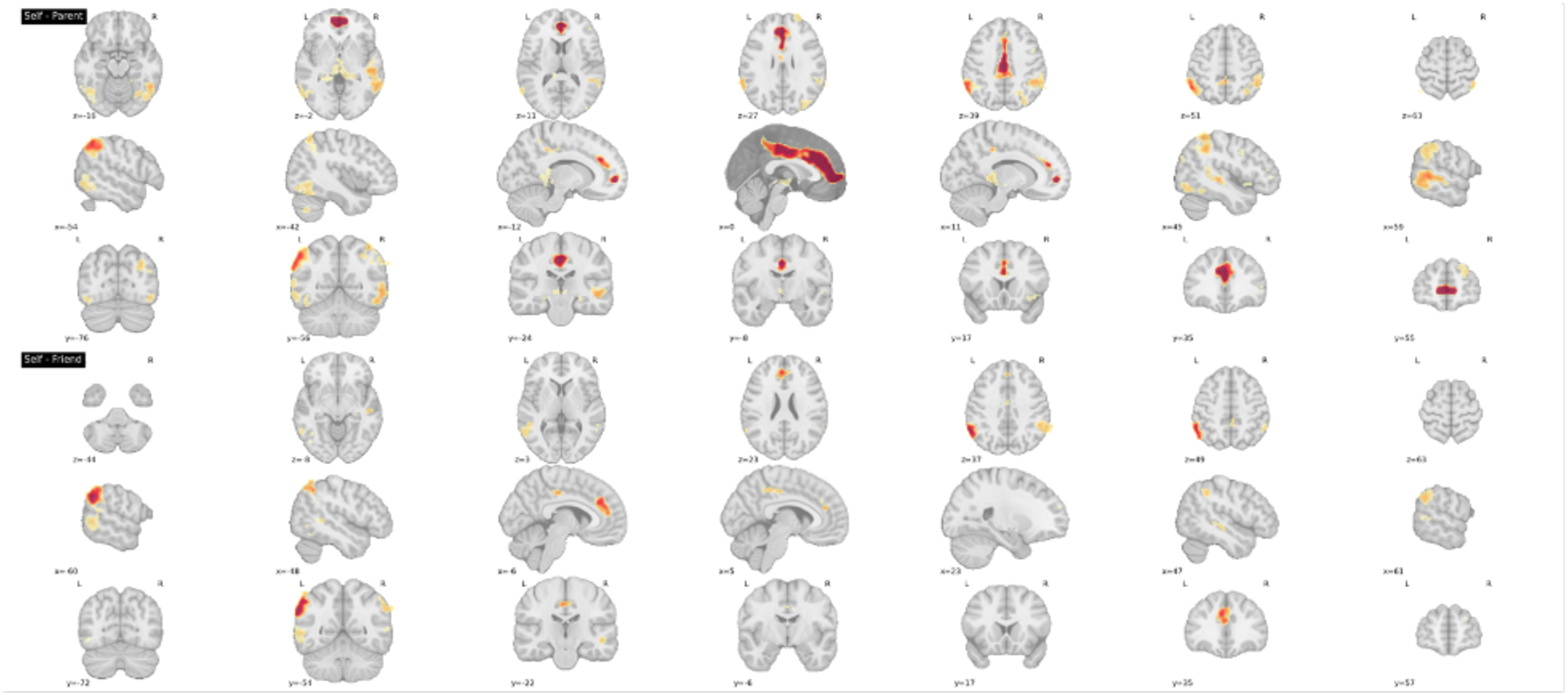
Univariate contrasts for the social preference and judgment task. *Note.* Group-level analyses were conducted via permutation test (10,000 iterations) using Nilearn’s non_parametric_inference function and TFCE. Negative log10 p-values are displayed as cluster intensity values. Clusters depict FWE-corrected *p-*values (*p* <.05).

**Figure 3-1.**
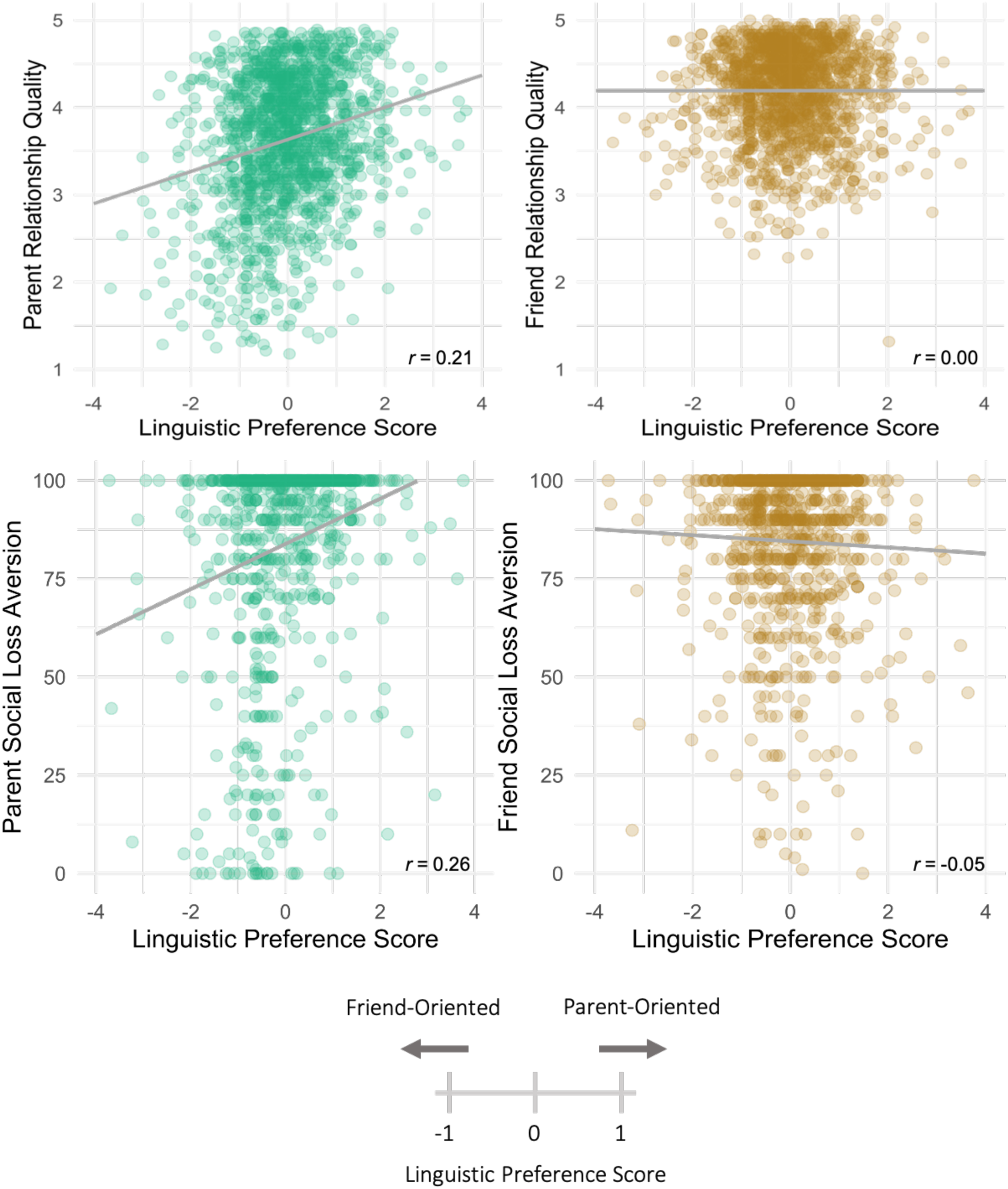
Linguistic preference scores strongly generalize to other relationship characteristics (relationship quality, social loss aversion) for parents and weakly generalize to social loss aversion for friends. *Note.* Linguistic preference scores refer to model implied social decision preferences between a parent and friend based on written text data; relatively greater values indicate a parent-over-friend preference, relatively lower values indicate a friend-over-parent preference. The Y-axis depicts relationship quality between participants and their nominated parent (left) and friend (right). Relationship quality was obtained with the self-administered IPPA. Social loss aversion is a one-shot item where participants are asked how upset they would be if they could no longer spend time with a given other. Linguistic preference scores were calculated with weights trained on the Word2Vec-based linguistic representations.

---

**Table 6-1.** Predicting social decision preferences with interactions between linguistic preference scores and neural dissimilarity scores (dmPFC, mPFC)

| Term | dmPFC – Monetary | mPFC – Monetary | dmPFC – Social | mPFC – Social |
| --- | --- | --- | --- | --- |
| Intercept | **-2.50 [-3.07, -1.93]** | **-2.44 [-3.03, -1.88]** | **-0.94 [-1.40, -0.52]** | **-0.89 [-1.33, -0.47]** |
| Condition | -0.03 [-0.55, 0.51] | -0.15 [-0.74, 0.38] | -0.28 [-0.78, 0.25] | -0.34 [-0.84, 0.16] |
| Reward ratio | -2.96 [-3.56, -2.40] | -2.95 [-3.52, -2.35] | -1.30 [-1.64, -0.97] | -1.29 [-1.61, -0.96] |
| Self-parent neural dissimilarity | 0.47 [0.00, 0.96] | 0.28 [-0.22, 0.76] | 0.03 [-0.35, 0.41] | 0.04 [-0.33, 0.43] |
| Self-friend neural dissimilarity | -0.20 [-0.66, 0.28] | 0.09 [-0.39, 0.56] | 0.05 [-0.35, 0.44] | 0.31 [-0.07, 0.68] |
| Linguistic preference score | 0.15 [-0.27, 0.61] | 0.21 [-0.25, 0.67] | -0.05 [-0.40, 0.30] | -0.07 [-0.42, 0.29] |
| Condition x Self-parent neural dissimilarity | -1.07 [-1.62, -0.51] | -0.30 [-0.87, 0.31] | -0.20 [-0.74, 0.37] | -0.05 [-0.59, 0.49] |
| Condition x Self-friend neural dissimilarity | -0.05 [-0.60, 0.48] | -0.11 [-0.70, 0.47] | -0.38 [-0.95, 0.20] | -0.60 [-1.17, -0.09] |
| Condition x Linguistic preference score | -0.50 [-1.01, 0.01] | -0.53 [-1.06, 0.05] | 0.19 [-0.33, 0.69] | 0.12 [-0.42, 0.62] |
| Self-parent neural dissimilarity x Linguistic preference score | 0.11 [-0.40, 0.62] | 0.18 [-0.35, 0.73] | 0.16 [-0.23, 0.55] | 0.02 [-0.39, 0.43] |
| Self-friend neural dissimilarity x Linguistic preference score | 0.08 [-0.47, 0.63] | -0.30 [-0.70, 0.47] | 0.12 [-0.33, 0.58] | -0.23 [-0.65, 0.21] |
| Condition x<br>Self-parent<br>neural<br>dissimilarity x<br>Linguistic<br>preference<br>score | -0.24 [-0.82, 0.35] | -0.11 [-0.77, 0.53] | -0.24 [-0.79, 0.32] | *-0.50 [-1.10, 0.08]* |
| Condition x<br>Self-friend<br>neural<br>dissimilarity x<br>Linguistic<br>preference<br>score | -0.64 [-1.07, 0.20] | 0.45 [-0.26, 1.12] | 0.08 [-0.56, 0.73] | -0.21 [-0.85, 0.39] |

*Note.* Coefficients are on a logit scale and reflect means from a posterior distribution. ‘Condition’ indicates when parent or friend outcomes were associated with the discounting or delay options and encodes social decision preferences (positive values indicate a parent-over-friend preference, negative values indicate a friend-over-parent preference). Reward ratio reflects the division of the non-discounting option over the discounting option. Self-parent and self-friend neural dissimilarities refer to the distance between multivoxel neural patterns for oneself and a parent or friend, extracted from the brain region listed in the column heading. Values in brackets reflect 89% highest density credible intervals drawn around the posterior distribution. Interactions reflect associations between neural dissimilarities, linguistic preferences, and/or social decision preferences. Coefficients where the HDI partially overlaps with ROPE (modest or moderate evidence depending on whether the interval contained 0) are italicized; coefficients where the HDI does not overlap with ROPE (strong evidence) are bolded and italicized.

**Table 6-2.**
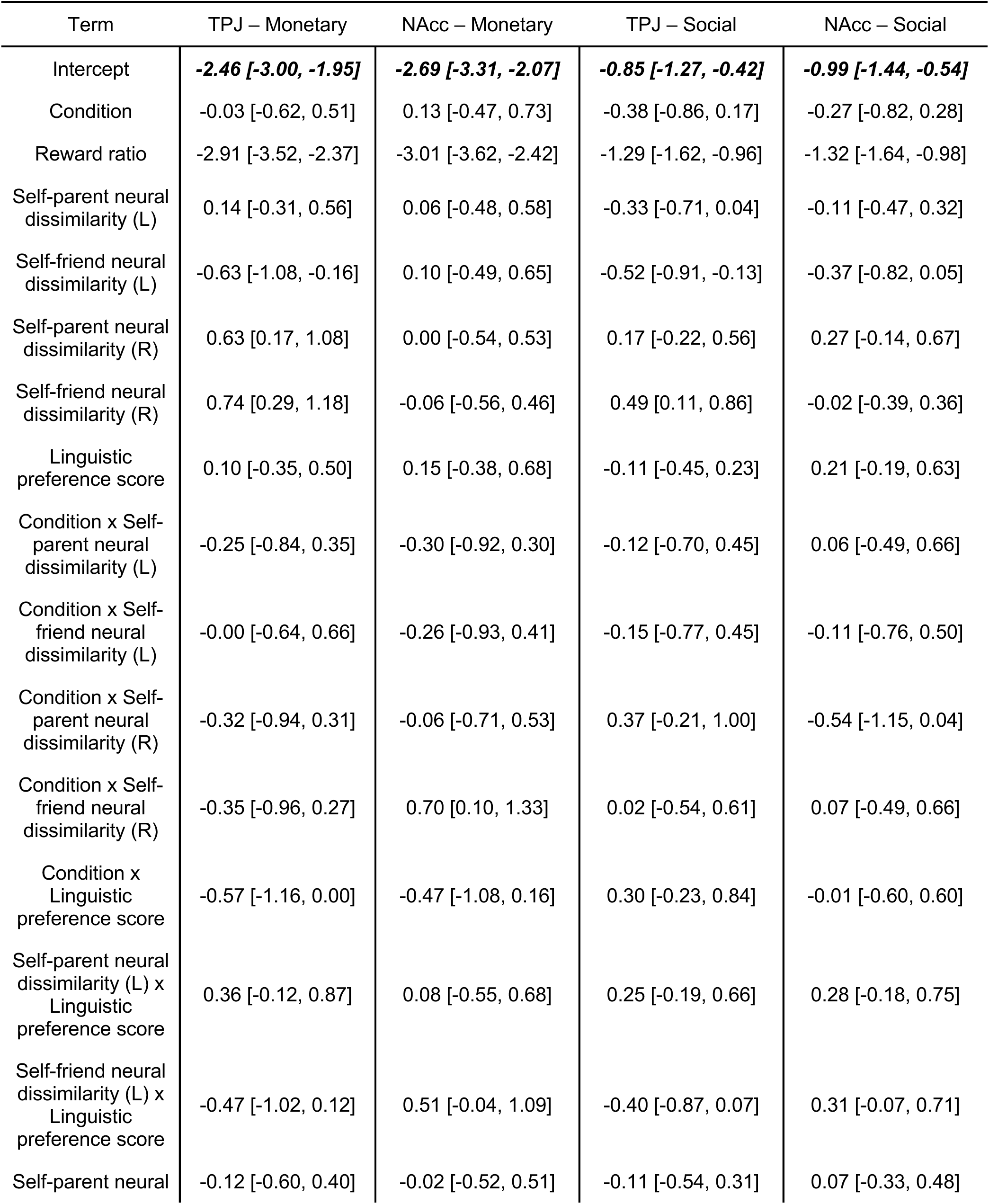
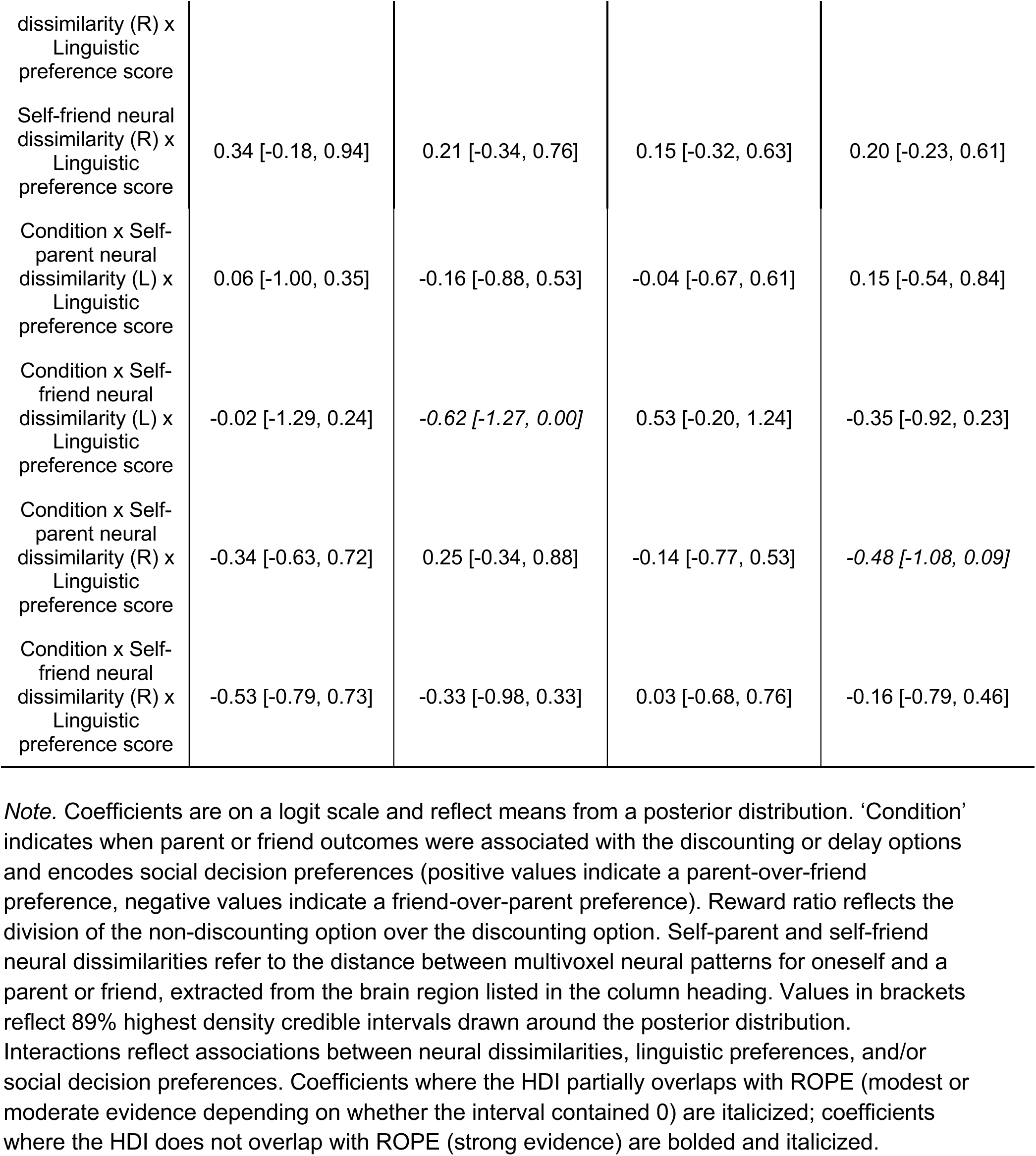
Predicting social decision preferences with interactions between linguistic preference scores and neural dissimilarity scores (TPJ, NAcc)

